# Human cytokine-induced memory-like NK cells preserve increased glycolysis but the glycolytic-dependence of their effector functions differ between stimuli

**DOI:** 10.1101/2020.08.20.258731

**Authors:** Iñigo Terrén, Ane Orrantia, Alba Mosteiro, Joana Vitallé, Olatz Zenarruzabeitia, Francisco Borrego

## Abstract

Natural Killer (NK) cells acquire memory-like properties following a brief stimulation with IL-12, IL-15 and IL-18. These IL-12/15/18-stimulated NK cells, also known as cytokine-induced memory-like (CIML) NK cells, have been revealed as a powerful tool in cancer immunotherapy due to their persistence in the host and their increased effector functions. Several studies have shown that NK cells modulate their metabolism in response to cytokine-stimulation and other stimuli, suggesting that there is a link between metabolism and cellular functions. In this paper, we have analyzed metabolic changes associated to IL-12/15/18-stimulation and the relevance of glycolytic pathway for NK cell effector functions. We have found that CIML NK cells are able to retain increased glycolytic machinery seven days after cytokine withdrawal. Furthermore, we found that glycolytic inhibition with 2-DG is stimuli-dependent and that differently affects to distinct effector functions. These findings may have implications in the design of NK cell-based cancer immunotherapies.

## INTRODUCTION

Natural Killer (NK) cells have the ability to directly kill tumor and virus-infected cells without prior sensitization. They can also release a wide variety of cytokines and chemokines that promote an adaptive immune response against target cells. Thus, NK cells represent a valuable tool in cancer immunotherapy, and several strategies have been proposed to exploit and improve their anti-tumor mechanisms in different cancers ^1–4^. Among them, a promising therapeutic approach is to stimulate NK cells with interleukins (ILs). For instance, IL-15 and superagonists, such as ALT-803, administration has been proven to be relatively safe and to effectively promote NK cell expansion in cancer patients ^5–8^. Alternatively, cells could be activated ex vivo prior to infusion. For example, IL-15-stimulated NK cells have been used in adoptive cell therapy protocols to treat different malignancies ^9,10^. Moreover, NK cells could be stimulated with combinations of ILs to enhance multiple effector functions. An example of a successful therapy following this strategy is the administration of IL-12/15/18-stimulated NK cells, also known as cytokine-induced memory-like (CIML) NK cells. These cells have been proven to be useful in mouse and rat models of several malignancies, including acute myeloid leukemia, T-cell acute lymphoblastic leukemia, multiple myeloma, lymphoma, melanoma, ovarian cancer and hepatocellular carcinoma ^11–17^. Regarding to humans, CIML NK cells have shown their safety and efficacy in the treatment of acute myeloid leukemia patients ^11^, and are currently being tested in several clinical trials (NCT01898793, NCT02782546, NCT03068819, NCT04024761, NCT04290546 and NCT04354025, from clinicaltrials.gov).

NK cells have been traditionally defined as innate lymphocytes, although this classification has become more challenging since adaptive NK cells were described in mice, non-human primates and humans ^18^. Unlike adaptive NK cells, CIML NK cells have not shown responses to specific antigens, although they share some features of immunological memory. These IL-12/15/18-preactivated NK cells were initially described in mice and were defined by their increased interferon gamma (IFNγ) production in response to a restimulation after a resting period ^19^. Moreover, these cells exhibited enhanced persistence in vivo and could be detected up to three months after adoptive transfer ^12,15^. Further studies revealed similar behavior of human IL-12/15/18-stimulated NK cells, showing an enhanced cytokine production following restimulation ^11,13,20–24^, and increased proliferation ^11,13,20^. After IL-12/15/18-stimulation, the phenotype of NK cells includes changes in the expression of activating and inhibitory receptors, and chemokine and cytokine receptors ^11,13,20,22,25–27^. Interestingly, it has been also demonstrated that cytokine-stimulation modifies the metabolic activity of NK cells ^28,29^, although this aspect has been poorly explored in human IL-12/15/18-stimulated NK cells.

Immune responses often involve changes in cellular metabolism, which is necessary to fulfill the different energetic demands of each cell function. Specifically related to NK cells, there is mounting evidence that the metabolic profile is modified during cell development, viral infection, and activation. This metabolic reprogramming is necessary to support processes such as IFNγ production ^30^, and includes changes in the activity of metabolic regulators, expression of nutrient transporters, and reconfiguration of the metabolic pathways ^31^. In this sense, it has been demonstrated that glucose metabolism is crucial for NK cell-mediated control of mouse cytomegalovirus infection ^32^. Glycolytic pathway is also essential in mouse NK cells for IFNγ and granzyme B production, although the mechanisms that regulate each function could be different ^33^. Certain pathologies such as obesity and cancer lead to altered metabolic activity, which has been linked to dysfunctional NK cells ^34–36^. Therefore, current knowledge suggests that there is a close relationship between NK cell metabolism and their effector functions.

Here, we have studied how human NK cell metabolism is modulated following IL-12/15/18-stimulation. Furthermore, considering that these preactivated NK cells have memory-like properties, and that they are being successfully used in clinical trials, we asked if the metabolic changes found immediately after the stimulation persisted for a long time. Since IL-15 and IL-2 have been previously used to promote NK cell survival and expansion, we tested the effect of both ILs after the preactivation with IL-12/15/18. Moreover, we have explored the different glycolytic requirements of several effector functions of IL-12/15/18-preactivated NK cells in response to different stimuli.

## METHODS

### Samples and cell culture

Blood samples (buffy coats) from adult healthy donors were collected through the Basque Biobank (biobancovasco.org). The Basque Biobank complies with the quality management, traceability and biosecurity, set out in the Spanish Law 14/2007 of Biomedical Research and in the Royal Decree 1716/2011. All subjects provided written and signed informed consent in accordance with the Declaration of Helsinki. The protocol was approved by the Basque Ethics Committee for Clinical Research (PI+INC-BIOEF 2014-02, PI+CES+INC-BIOEF 2017-03 and PI2014079). Fresh peripheral blood mononuclear cells (PBMCs) were obtained from buffy coats by Ficoll Paque Plus (GE Healthcare) density gradient centrifugation. NK cells were isolated from PBMCs by negative depletion with human NK cell Isolation Kit (Miltenyi Biotec). Cells were cultured in RPMI 1640 medium supplemented with GlutaMAX (Gibco), 10% heat-inactivated Fetal Bovine Serum (FBS) (HyClone), 1% non-essential amino acids (Gibco), 1% Sodium Pyruvate (Gibco) and 1% Penicillin-Streptomycin (Gibco). PBMCs or purified NK cells were stimulated for 16-18 hours with 10 ng/mL rhIL-12 (Miltenyi Biotec), 100 ng/mL rhIL-15 (Miltenyi Biotec) and 50 ng/mL rhIL-18 (MBL International Corporation), or cultured in media alone for the same time. Cells were then washed three times and cultured with 1 ng/mL rhIL-15 or 20 IU/mL rhIL-2 (Miltenyi Biotec). After 3 days, media and cytokines (IL-15 or IL-2) were replaced, and cells were cultured for 4 more days. The time-points after the 16-18 hours stimulation, and after seven days of culture, would be hereinafter referred to as Day 0 and Day 7, respectively (**Figure S1**). The K562 cell line was cultured in the same media than NK cells supplemented with 5 µg/mL of Plasmocin (InvivoGen) and was routinely tested for mycoplasma infection with Venor GeM Classic detection kit (Minerva Biolabs).

### Flow cytometry

Extracellular staining was performed by incubating cells for 30 minutes on ice with the following fluorochrome-conjugated mouse anti-human antibodies: BV421 anti-CD71 (M-A712), BV510 anti-CD3 (UCHT1), BV510 anti-CD14 (MφP9) and PE anti-CD98 (UM7F8) from BD Biosciences; and PE-Vio770 anti-CD56 (REA196) from Miltenyi Biotec. Cells were then washed with PBS containing 2.5% Bovine Serum Albumin (BSA) (Sigma-Aldrich), and, for intracellular staining, cells were fixed and permeabilized with Fixation/Permeabilization solution (BD Biosciences), following manufacturer’s protocol. Next, intracellular staining was performed by incubating cells for 30 minutes on ice with the following fluorochrome conjugated mouse anti-human antibodies: BV421 anti-IFNγ (B27) and FITC anti-MIP-1β (D21-1351) from BD Biosciences; and APC anti-TNF (Mab11) from BioLegend. The following fluorochrome conjugated rabbit anti-human antibodies were also used in the intracellular staining: FITC anti-GLUT3 (polyclonal) and PE anti-GLUT1 (EPR3915) from Abcam. Brilliant Stain Buffer (BD Biosciences) was used during the simultaneous incubation with BV421 and BV510 dyes. Cell viability was determined by staining cells for 30 minutes on ice with LIVE/DEAD Fixable Near-IR Dead Cell Stain Kit (Invitrogen) prior to extracellular staining. Mitochondrial mass was measured by incubating cells with 150 nM MitoTracker Green FM (Invitrogen) for 25 min at 37°C after the extracellular staining. Glucose uptake assay was performed by incubating cells with 50 µM 2-NBDG (2-(N-(7-Nitrobenz-2-oxa-1,3-diazol-4-yl)Amino)-2-Deoxyglucose) (Invitrogen) for 90 min at 37°C prior to cell staining. NK cells were gated within viable lymphocytes as CD3-CD14-CD56+ cells (**Figure S2**). Flow cytometry was performed with MACSQuant Analyzer 10 (Miltenyi Biotec). Data was analyzed with FlowLogic v.7.3 software.

### Extracellular flux assay

Purified NK cells were washed and resuspended in Seahorse XF DMEM Medium (pH 7.4, Agilent Technologies) supplemented with 10 mM glucose, 2 mM glutamine and 1 mM pyruvate. 96 well microplates were treated with Cell-Tak (Corning) and cells were plated into six replicates at 200,000 cells/well. Extracellular acidification rate (ECAR) and oxygen consumption rate (OCR) were determined using a Seahorse XFe96 Extracellular Flux Analyzer (Agilent Technologies). Seahorse XF Cell Mito Stress Test (Agilent Technologies) was used following manufacturer’s indications, using 1, 1 and 0.5 µM of oligomycin, FCCP and rotenone/antimycin A, respectively. Data was analyzed with Seahorse Wave Desktop software v.2.6 (Agilent Technologies).

### Functional assay

NK cells were washed and plated at 200,000 cell/well (10^6^ cell/mL) in 96 U-bottom well plates in the presence and absence of 50 mM 2-Deoxy-D-glucose (2-DG) (Sigma-Aldrich). To stimulate them, K562 target cells at 1:1 effector:target (E:T) ratio, or IL-12, IL-15 and IL-18 (10, 100 and 50 ng/mL, respectively) were added. Mouse anti-human PE anti-CD107a mAb (REA792, from Miltenyi Biotec) was added and cells were cultured at 37°C for one hour. Then, GolgiPlug and GolgiStop (brefeldin A and monensin, respectively) protein transport inhibitors (BD Biosciences) were added, following manufacturer’s recommendations, and cells were cultured for additional 6 hours at 37°C. Plates were pulse centrifuged at 200 g for 1 minute prior to each incubation period. Cells were then collected, stored overnight at 4°C, and stained and analyzed by flow cytometry the next day. The percentage of specific response was calculated as the percentage of stimulated cells that are positive for a specific function minus the percentage of non-stimulated cells that are positive for the same function.

### Cytotoxicity assay

The assay was performed following a previously described protocol ^20^. Briefly, K562 target cells were incubated for 30 minutes at 37°C in the presence of 15 µM calcein-AM (Invitrogen). PBMCs were washed and plated into three replicates in 96 U-bottom well plates with labeled target cells at different E:T ratios, with 5,000 target cells per well. Plates were pulse centrifuged at 200 g for 1 minute and then cultured for 3 hours at 37°C in the presence and absence of 50 mM 2-DG. Then, plates were centrifuged and 75 µL of the supernatant were transferred to black 96 well plates to measure calcein-AM release with a Varioskan Flash fluorimeter (Thermo Fisher Scientific), with a configuration of excitation/emission of 485/518 nm. Mean of the triplicates of each condition was calculated for the analysis. Medium fluorescence was calculated without cells, in the presence and absence of 2-DG. Spontaneous release was measured by culturing target cells without PBMCs, in the presence and absence of 2-DG. Maximum release was measured by culturing labeled target cells with 2% Triton X-100 (Sigma-Aldrich), in the presence and absence of 2-DG. Triton fluorescence was measured with media plus 2% Triton X-100 without cells, in the presence and absence of 2-DG. The percentage of specific lysis was calculated with the following formula:

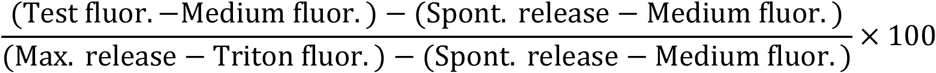

### Statistical analysis and data representation

GraphPad Prism v.8.4 was used for graphical representation and statistical analysis. Data was represented showing mean ± standard error of the mean (SEM). Non-parametric Wilcoxon matched-pairs signed rank test were used to determine significant differences. ns: non-significant, *p<0.05, **p<0.01.

## RESULTS

### IL-12/15/18-preactivated NK cells exhibit increased expression of nutrient transporters

Previous studies have shown that NK cells can modify the expression of nutrient transporters following cytokine-stimulation ^30,37–43^. In accordance, we have found that NK cells increase the expression of the transferrin receptor CD71, amino acid transporter CD98, and glucose transporters GLUT1 and GLUT3, following stimulation with IL-12, IL-15 and IL-18 (hereinafter referred to as IL-12/15/18-preactivated or CIML NK cells) for 16-18 hours, i.e. at day 0 (**Figure 1, left column**).

**Figure 1.**
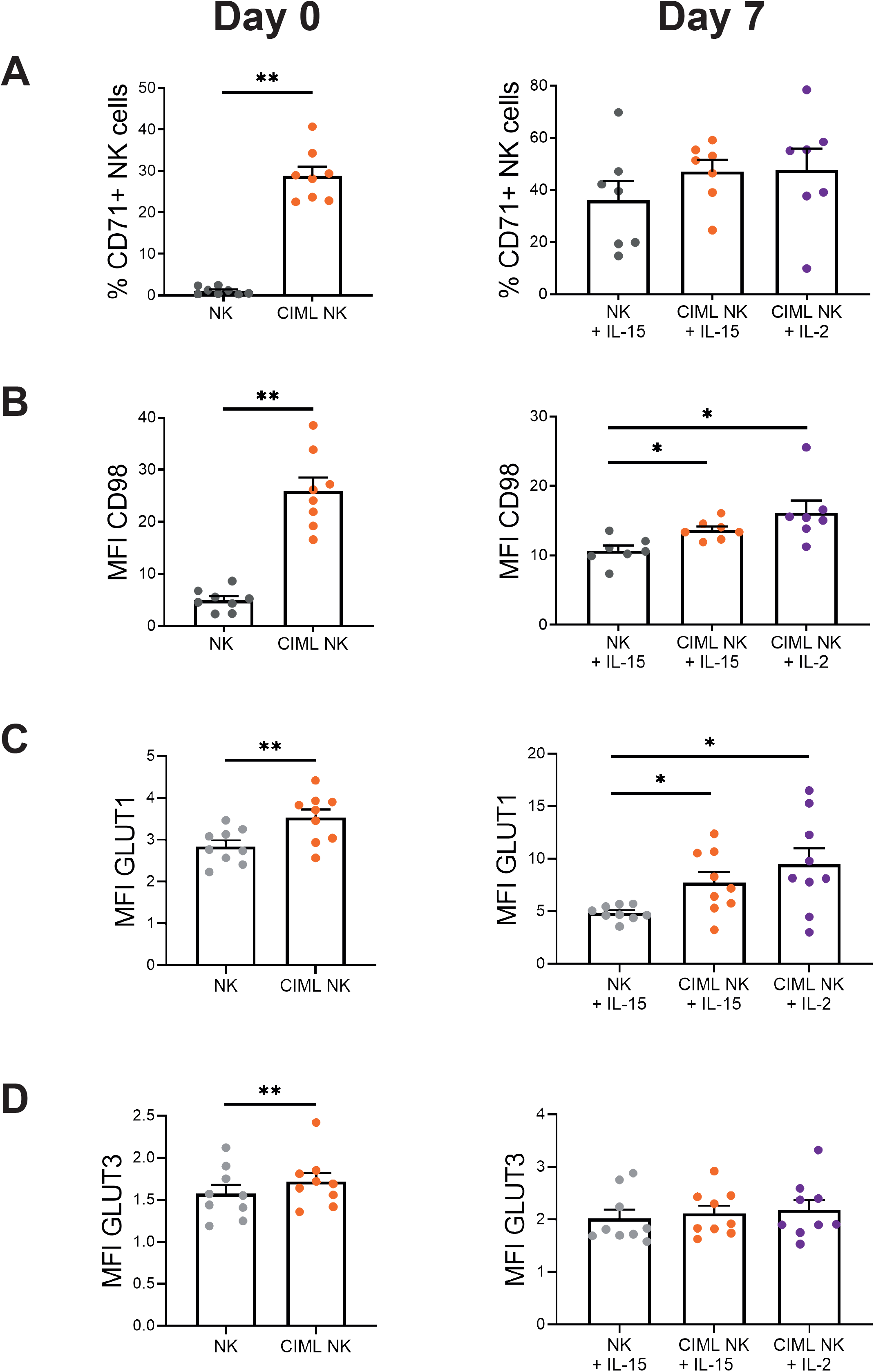
Expression of nutrient transporters in human cytokine-induced memory-like (CIML) NK cells. Bar charts representing (**A**) the percentage of control NK cells (NK) and CIML NK cells expressing the transferrin receptor CD71 and expression levels of (**B**) the amino acid transporter CD98 and glucose transporters (**C**) GLUT1 and (**D**) GLUT3, measured as median fluorescence intensity (MFI). Graphs show data of cells after 16-18 hours of IL-12/15/18-stimulation (Day 0, left column) and after 7 days of culture with IL-15 (NK+IL-15 or CIML NK+IL-15) or IL-2 (CIML NK+IL-2) (Day 7, right column). Means ± SEM are depicted. Statistical analyses were performed using Wilcoxon matched-pairs signed rank test. Each dot represents an independent experiment from a different donor (n = 6-9). *p<0.05, **p<0.01.

We next asked if the increased expression of nutrient transporters returned to resting levels after cytokine withdrawal, or if it was maintained when cells returned to a less activated state. We cultured control and CIML NK cells with low doses of IL-15 (1 ng/mL) for seven days, as others have previously done ^11,24^, to support their survival without inducing a strong activation. By doing so, we expected that cells could return to a resting situation. Furthermore, IL-12/15/18-preactivated NK cells have an increased expression of alpha chain of the IL-2 receptor (IL-2Rα, also known as CD25), which conforms the high-affinity IL-2 receptor ^11–13,20,22,25,26^. This feature allows CIML NK cells to proliferate in response to low doses of IL-2, as we and others have previously described ^12,20,22^. Therefore, we washed control and CIML NK cells, cultured them for a period of seven days with low concentrations of IL-15 or IL-2 (**Figure S1**), and then analyzed again the expression of nutrient transporters. Our results showed that CIML NK cells express higher levels of CD98 and GLUT1 than control NK cells at day 7 (**Figure 1, right column**). On the other hand, no significant differences were found in the expression of CD71 and GLUT3 between control and CIML NK cells (**Figure 1, right column**). Of note, CD71 is not (or very low) expressed in resting NK cells, but culturing them for seven days with low doses of IL-15 was enough to induce CD71 expression (**Figure 1 and Figure S3**). Altogether, these results show that NK cells increase the expression of nutrient transporters following stimulation with IL-12, IL-15 and IL-18, and that CIML NK cells preserve an elevated expression of amino acid and glucose transporters when they are maintained in culture with very low concentrations of IL-2 or IL-15.

### IL-12/15/18-preactivated NK cells retain increased glycolytic activity

Considering that IL-12/15/18-preactivated NK cells have an increased GLUT1 expression, and that glycolysis supports multiple NK cell functions ^31,36^, we next analyzed changes in this metabolic pathway. In accordance with the expression of GLUT1, glucose uptake assay with the glucose analogue 2-NBDG revealed that the consumption of this substrate is higher in CIML NK cells at both time-points, i.e. at day 0 and day 7 (**Figure 2A**). We further studied glycolytic activity of NK cells with the extracellular flux analyzer Seahorse XF. Our results showed that NK cells tended to increase the glycolytic rate, measured as extracellular acidification rate (ECAR), following stimulation with IL-12, IL-15 and IL-18 (**Figure 2B, left panel**). These results support the hypothesis that NK cells increase glycolytic activity following stimulation ^31^. Interestingly, CIML NK cells tend to show a higher glycolytic rate than control NK cells at day 7, especially when cultured with low doses of IL-2 (**Figure 2B, right panel**). Oxidative phosphorylation (OXPHOS) levels, measured as oxygen consumption rate (OCR), also tended to increase in NK cells after the initial cytokine-stimulation. However, the differences on OXPHOS levels between control and IL-12/15/18-preactivated NK cells were negligible after seven days of culture (**Figure 2C**). Then, we calculated the ratio between OXPHOS and glycolytic rates, which has been previously used to detect shifts in fuel utilization ^44^. Our results showed that CIML NK cells tended to have a lower OCR:ECAR ratio, although the difference is better observed after seven days, indicating that CIML NK cells undergo a metabolic switch towards glycolysis that is not stopped following cytokine withdrawal (**Figure 2D**). Altogether, these results support the hypothesis that IL-12/15/18-preactivated NK cells increase their glycolytic machinery, and that they can retain it for at least seven days when cultured with low concentrations of IL-15 or IL-2.

**Figure 2.**
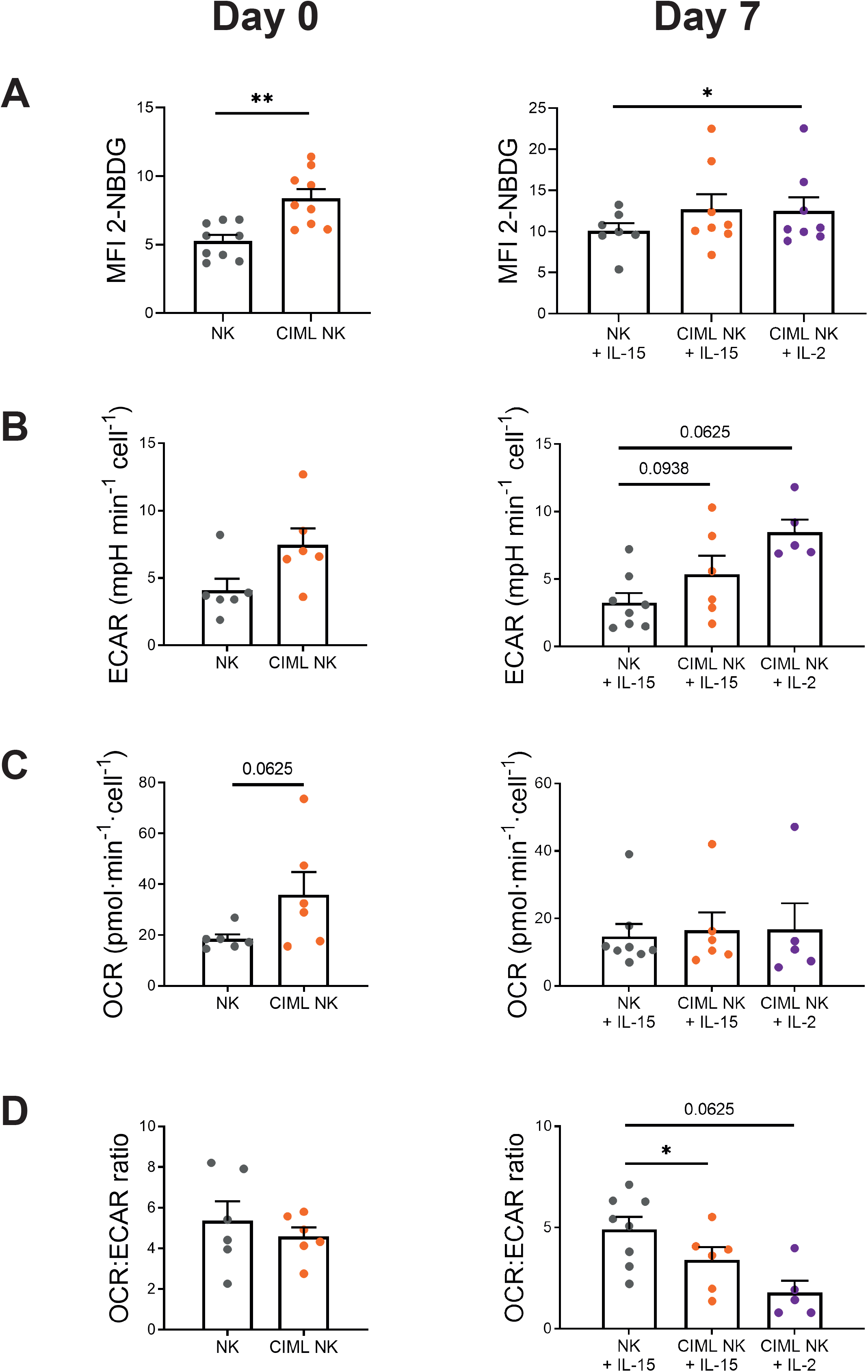
Metabolic reprogramming of human cytokine-induced memory-like (CIML) NK cells. Bar charts representing (**A**) glucose uptake, measured as median fluorescence intensity (MFI) of the glucose analogue 2-NBDG. Bar charts representing (**B**) glycolytic rate, measured as extracellular acidification rate (ECAR); (**C**) OXPHOS rate, measured as oxygen consumption rate (OCR); and (**D**) OCR:ECAR ratio. Graphs show data of control NK cells (NK) and IL-12/15/18-stimulated NK cells (CIML NK) after 16-18 hours (Day 0, left column), and after 7 days of culture with IL-15 (NK+IL-15 or CIML NK+IL-15) or IL-2 (CIML NK+IL-2) (Day 7, right column). Means ± SEM are depicted. Statistical analyses were performed using Wilcoxon matched-pairs signed rank test. Each dot represents an independent experiment from a different donor (n = 5-9). *p<0.05, **p<0.01.

### Increased mitochondrial activity is not sustained for long periods

OXPHOS activity tended to increase following IL-12/15/18-stimulation, but it returned to basal levels after a week in the absence of strong stimuli (**Figure 2C**). We decided to further explore the mitochondrial activity by analyzing the spare respiratory capacity (SRC), defined as the difference between the maximal and basal respiration (**Figure S4**). In accordance with OXPHOS levels, NK cells showed increased SRC after the initial IL-12/15/18-stimulation, but no significant differences were found at day 7 (**Figure 3A and 3B**). Similarly, levels of ATP-linked respiration and non-mitochondrial oxygen consumption were significantly different after IL-12/15/18-stimulation, but similar after 7 days of culture (**Figure S5**). To examine if the differences found in the mitochondrial activity were due to changes in the cellular mitochondrial content, we next analyzed mitochondrial mass using MitoTracker staining. In contrast to other publications where 18 hours of IL-2/12-stimulation increased mitochondrial mass ^38^, we did not found differences in the mitochondrial mass of control and IL-12/15/18-stimulated NK cells at day 0 (**Figure 3C, left panel**). Surprisingly, we found significant differences between control and CIML NK cells after 7 days of culture (**Figure 3C, right panel**). We dismissed the hypothesis that increased mitochondrial mass was due to a higher cellular size since we found that CIML NK cells were larger than control NK cells at both day 0 and day 7 (**Figure S6A**). These data suggest that IL-12/15/18-stimulation does not rapidly induce mitochondrial biogenesis of NK cells. Instead, mitochondrial mass is progressively increased during the following seven days of culture (**Figure S6B**). Also, our data demonstrate that increased mitochondrial mass does not necessarily correlate with augmented OXPHOS activity, since IL-12/15/18-preactivated NK cells show higher mitochondrial mass than control NK cells at day 7 (**Figure 3C, right panel**) but similar OXPHOS and SRC rates (**Figure 2C and 3B**).

**Figure 3.**
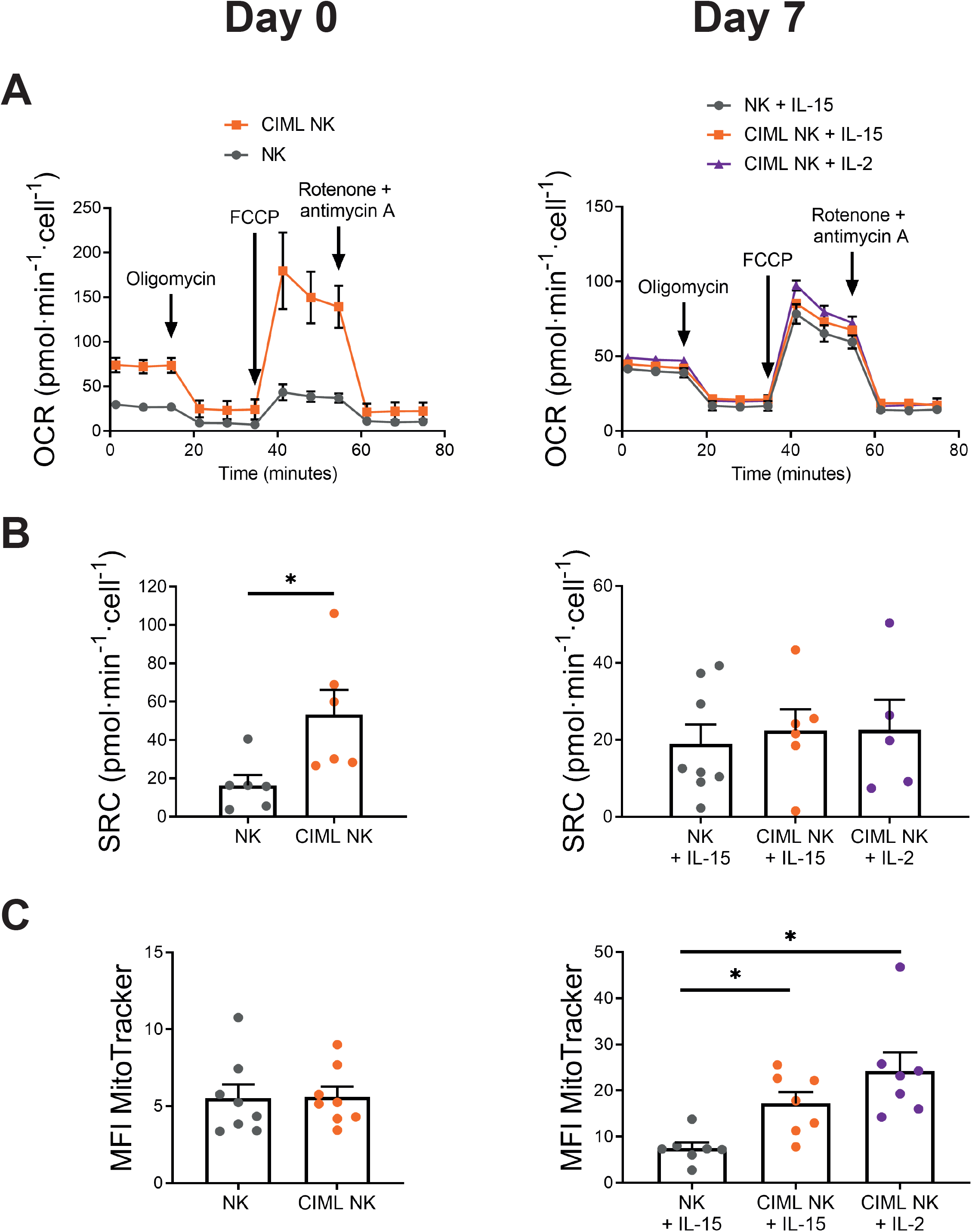
Mitochondrial activity and mass of human cytokine-induced memory-like (CIML) NK cells. (**A**) Representative experiment of Seahorse XF analyzer with the Mito Stress Test kit. Means ± standard deviation are depicted. (**B**) Bar charts representing spare respiratory capacity (SRC), measured as the difference between maximal and basal mitochondrial respirations. (**C**) Bar charts representing mitochondrial mass, measured as the median fluorescence intensity (MFI) of MitoTracker Green. Graphs show data of control NK cells (NK) and IL-12/15/18-stimulated NK cells (CIML NK) after 16-18 hours (Day 0, left column), and after 7 days of culture with IL-15 (NK+IL-15 or CIML NK+IL-15) or IL-2 (CIML NK+IL-2) (Day 7, right column). Means ± SEM are depicted in bar graphs. Statistical analyses were performed using Wilcoxon matched-pairs signed rank test. Each dot represents an independent experiment from a different donor (n = 5-8). *p<0.05.

### Glycolytic inhibition differently affects to distinct NK cell effector functions and is stimuli-dependent

Elevated rates of glycolysis are required for maximal IFNγ production in mouse NK cells ^33,43^. Similarly, inhibiting the glycolytic pathway by using galactose as an alternative carbon fuel decreased IFNγ production of human IL-12/15-stimulated CD56^bright^ NK cells, but not of IL-2-stimulated NK cells ^30^. Interestingly, some authors have reported that inhibiting glycolysis had minimal inhibitory effects on cytokine-stimulated NK cell degranulation ^30,32^, suggesting that sensitivity to glycolytic inhibition may be different among distinct effector functions, but that also depends on the stimuli. Considering that IL-12/15/18-preactivated NK cells have elevated rates of glycolysis, we decided to explore the effect of the glycolysis inhibitor 2-DG on their different effector functions using distinct stimuli. Once 2-DG is taken up into cells, it is phosphorylated by hexokinase to 2-deoxy-D-glucose-6-phosphate, which cannot be further metabolized, resulting in its cellular accumulation and thus inducing competitive and non-competitive inhibition of glucose-6-phosphate isomerase and hexokinase, respectively ^45^. Hence, CIML NK cells were co-incubated with K562 target cells for 7 hours in the presence and absence of 2-DG. Cytokine and chemokine production (IFNγ, TNF and MIP-1β) and degranulation (CD107a) were measured, and the percentage of specific responding cells was calculated by subtracting the percentage of positive cells, for each studied function, in the absence of stimulus. This calculation was especially relevant after the 16-18 hours of cytokine-stimulation, which increased cytokine and chemokine production, and degranulation, of CIML NK cells (**Figure S7**). Results showed that all the studied effector functions of both, control and IL-12/15/18-preactivated NK cells, were reduced in the presence of 2-DG. However, IFNγ and MIP-1β production was drastically reduced, while TNF production and degranulation were only partially inhibited (**Figure 4A**). We further studied the effect of glycolytic inhibition over NK cell effector functions by analyzing their cytotoxic activity against K562 target cells in the presence of 2-DG. Interestingly, our results showed that the specific lysis of K562 cells was reduced, but not completely inhibited. Notably, CIML NK cells showed higher cytotoxic activity than control NK cells, even in the presence of 2-DG (**Figure 4B**).

**Figure 4.**
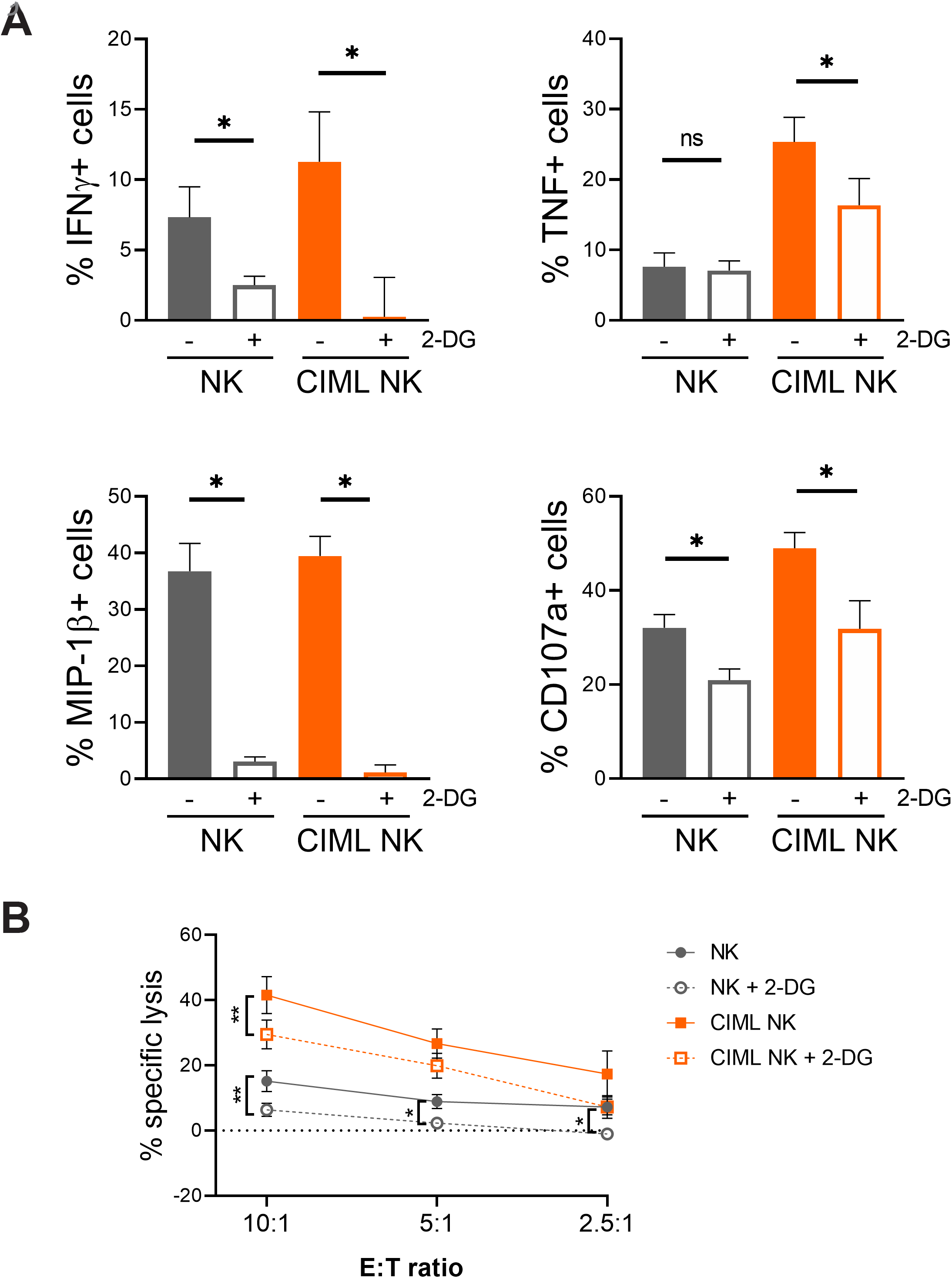
2-DG-induced inhibition of specific cytokine/chemokine production, degranulation and cytotoxic activity of human cytokine-induced memory-like (CIML) NK cells at day 0. (**A**) Control (NK) and CIML NK cells were co-cultured with K562 target cells (E:T ratio = 1:1) for 7 hours in the presence and absence of 50 mM 2-DG. Bar graphs showing the specific percentage of cells that produce IFNγ, TNF and MIP-1β, or degranulate (CD107a), in the presence and absence of 2-DG. Specific percentage of positive cells was calculated by subtracting the percentage of positive cells for each measured function in the absence of stimuli. Means ± SEM are depicted. Statistical analyses were performed using Wilcoxon matched-pairs signed rank test. n = 7. (**B**) Control (NK) and CIML NK cells were co-cultured with calcein-AM-labeled K562 target cells at different E:T ratios for 3 hours in the presence and absence of 50 mM 2-DG. Percentage of specific lysis was measured based on calcein-AM release. Means ± SEM are depicted. Statistical analyses were performed using Wilcoxon matched-pairs signed rank test. n = 7-8. ns = non-significant, *p<0.05, **p<0.01.

As previously described, we also cultured NK cells for 7 days with low concentrations of IL-15 or IL-2, and then analyzed their effector functions. Unlike previous experiments in which CIML NK cells were highly activated due to cytokine-stimulation for 16-18 hours, after seven days of culture period with low cytokine concentrations, control and IL-12/15/18-preactivated NK cells showed no differences in cytokine and chemokine production, and degranulation, in the absence of additional stimuli (**Figure S8**). Considering that CIML NK cells were characterized by a higher IFNγ production after cytokine-restimulation ^19^, we decided to stimulate CIML NK cells either with K562 target cells or with a mixture of IL-12, IL-15 and IL-18. Our results showed that IFNγ production was inhibited by 2-DG in both control and CIML NK cells, although there was a higher percentage of responding cells in the latter group (**Figure 5**). MIP-1β production was again strongly inhibited in both control and CIML NK cells (**Figure 5**). Interestingly, TNF production was inhibited when NK cells were stimulated with ILs, but not when they were co-cultured with K562 target cells (**Figure 5**). Of note, it should be considered that TNF production in response to cytokine-stimulation was very low in comparison with K562-stimulation. Similarly, degranulation of NK cells tended to be more inhibited by 2-DG when cells were stimulated with ILs, in comparison with K562-stimulation, although no significant differences were observed (**Figure 5**). Moreover, we also analyzed cytotoxic activity of control and CIML NK cells against K562 cells at day 7. We found that their cytotoxicity was reduced but not completely inhibited by 2-DG in all the studied E:T ratios (**Figure 6**).

**Figure 5.**
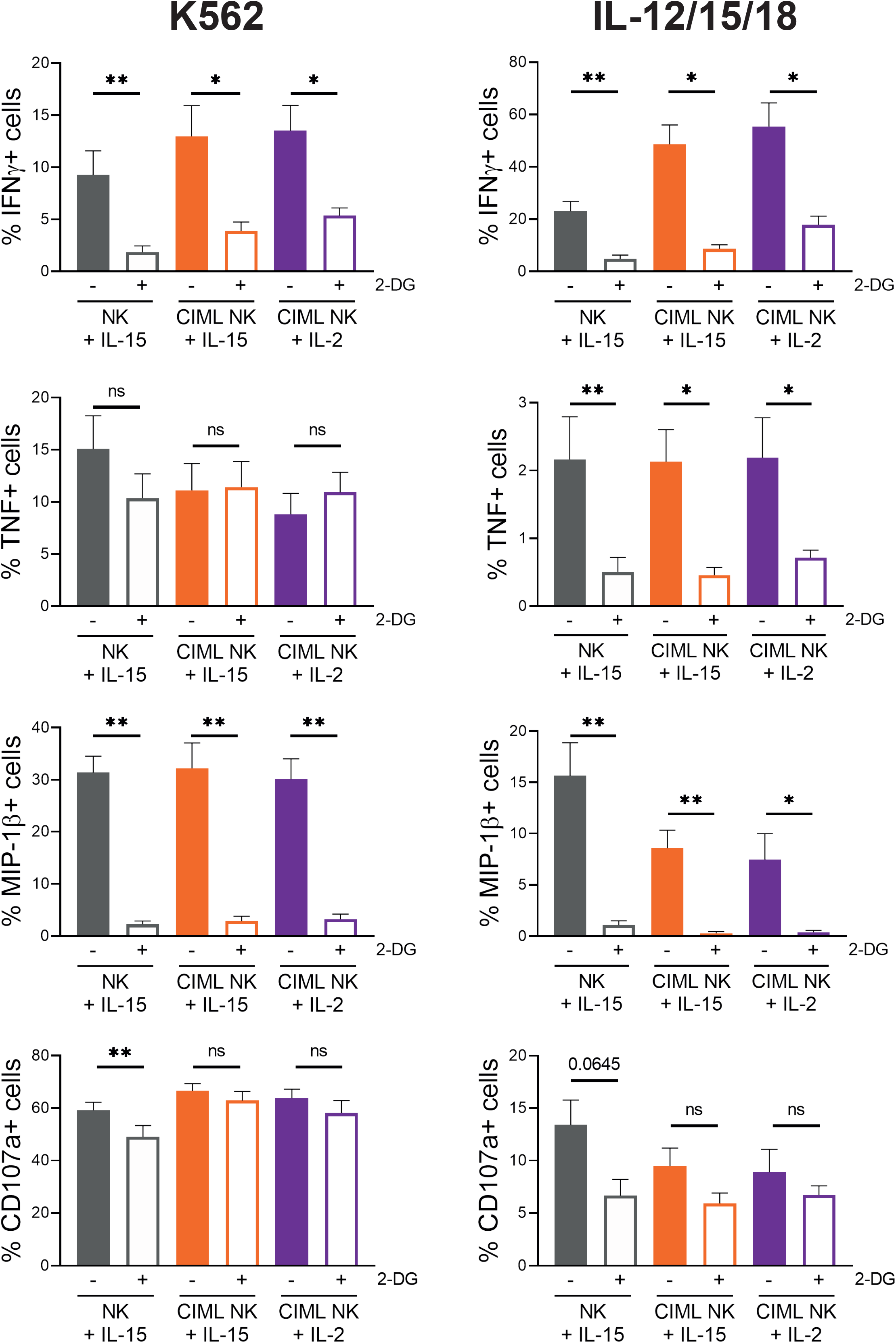
2-DG-induced inhibition of specific cytokine/chemokine production and degranulation of human cytokine-induced memory-like (CIML) NK cells at day 7. Control (NK+IL-15) and CIML NK cells cultured for seven days with IL-15 (CIML NK+IL-15) or IL-2 (CIML NK+IL-2), were co-cultured with K562 target cells (E:T ratio = 1:1) or restimulated with IL-12, IL-15 and IL-18 (10, 100 and 50 ng/mL, respectively) for 7 hours in the presence and absence of 50 mM 2-DG. Bar graphs showing the specific percentage of cells that produce IFNγ, TNF and MIP-1β, or degranulate (CD107a), in the presence and absence of 2-DG. Specific percentage of positive cells was calculated by subtracting the percentage of positive cells for each measured function in the absence of stimuli. Means ± SEM are depicted. Statistical analyses were performed using Wilcoxon matched-pairs signed rank test. n = 7-9. ns = non-significant, *p<0.05, **p<0.01.

**Figure 6.**
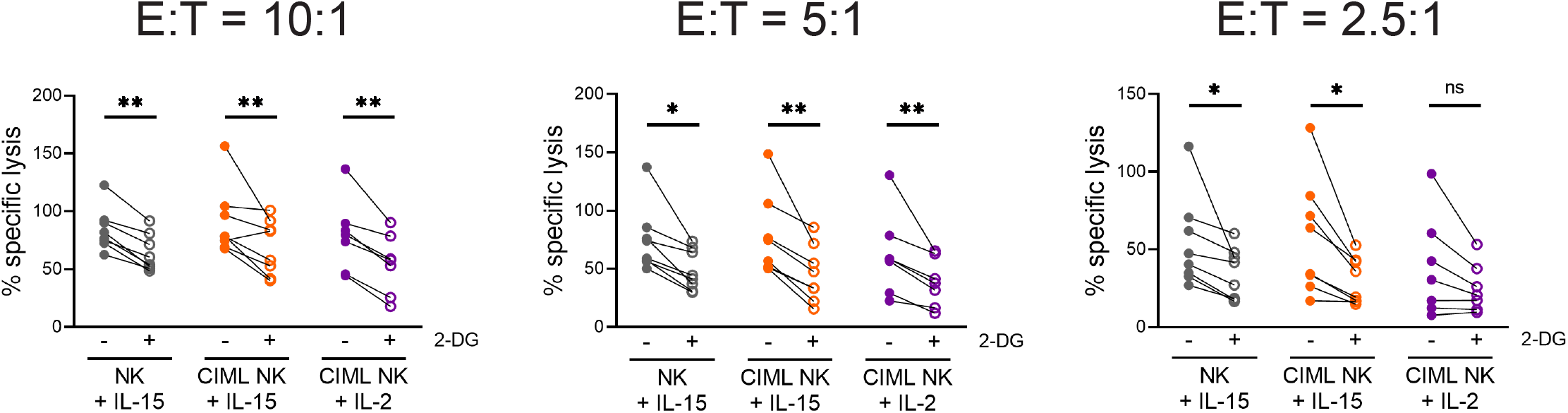
2-DG-induced inhibition of cytotoxic activity of human cytokine-induced memory-like (CIML) NK cells at day 7. Control (NK+IL-15) and CIML NK cells cultured for seven days with IL-15 (CIML NK+IL-15) or IL-2 (CIML NK+IL-2), were co-cultured with calcein-AM-labeled K562 target cells at different E:T ratios for 3 hours in the presence and absence of 50 mM 2-DG. Percentage of specific lysis was measured based on calcein-AM release. Means ± SEM are depicted. Statistical analyses were performed using Wilcoxon matched-pairs signed rank test. n = 7-8. ns = non-significant, *p<0.05, **p<0.01.

Altogether, our results showed that effector functions of NK cells were differently affected by 2-DG. Our data suggest that IFNγ and MIP-1β production are very sensitive to glycolysis inhibition, while TNF production, degranulation and cytotoxic activity are reduced but not entirely inhibited when glycolysis is blocked. Moreover, our data suggests that 2-DG-mediated glycolysis inhibition could be dependent on the stimuli, i.e. target cells vs. cytokine-stimulation.

## DISCUSSION

IL-12/15/18-preactivated NK cells have emerged as a powerful tool to treat leukemia ^11^, and their potential is also being explored in therapies against other malignancies ^13–15,17^. One of the characteristics of CIML NK cells that exemplify an advantage in cancer treatment is their persistence in the host. Ni et al. demonstrated that IL-12/15/18-preactivated NK cells were detectable 3 months after adoptive transfer in RMA-S tumor-bearing mice ^12^. The long-term benefits of cytokine-activated NK cells have been proven in a mouse model of multiple myeloma, in which IL-12/15/18-stimulated NK cells were able to reduce tumor burden in bone marrow after 7 days, while the treatment with IL-15-stimulated NK cells failed ^15^. Similarly, in a xenogeneic mouse model of ovarian cancer, tumor suppression was maintained after 20 days only in mice treated with CIML NK cells, but not with IL-15-stimulated NK cells ^13^. Considering the long-term effects of these cells, we decided to analyze the metabolic features of IL-12/15/18-preactivated NK cells both before and after a culture period of 7 days. Current therapeutic approaches include the in vivo administration of IL-15, IL-2 or modified versions of these cytokines after cell infusion to promote survival and expansion of CIML NK cells in the host (NCT01898793, NCT02782546, NCT04024761, NCT04290546 and NCT04354025, from clinicaltrials.gov). In our experimental settings, we sought to induce NK cell survival while avoiding excessive stimulation, so we cultured both control and CIML NK cells with very low doses of IL-15 for one week, as others have previously done ^11,24^. Additionally, we tried to mimic the conditions in which infused CIML NK cells in clinical trials are helped to expand ^11^, by culturing them with IL-2. Although we used low doses of IL-2 (20 IU/mL), we have previously described that this concentration is enough to promote proliferation of IL-12/15/18-preactivated NK cells ^20^. Therefore, our experimental design allowed us to in vitro study the metabolism and effector functions of resting and proliferating CIML NK cells, in a somehow comparable manner as it may be happening after their administration in patients.

We first decided to analyze the expression of nutrient transporters. Our data is in accordance with results published by others, confirming that the expression of nutrient transporters in NK cells can be upregulated in response to different stimuli ^30,33,37–43,46^. Interestingly, our results showed that 7 days after cytokine-preactivation, CIML NK cells retain increased expression of CD98 and GLUT1, although the relevance of these transporters in the tumor context is still unknown. Inhibition of CD98 activity with D-phenylalanine has been found to completely abolish NKG2D-induced IFNγ production and degranulation of NK cells ^42^. Similarly, inhibition of amino acid transport through CD98/LAT1 complex with 2-aminobicyclo-(2,2,1)-heptane-2-carboxylic acid (BCH) inhibited IFNγ and granzyme B production of IL-2/18-and IL-2/12-stimulated NK cells ^38,39^. Overall, these reports showed that blocking CD98 activity limits NK cells effector functions, but to our knowledge, an elevated expression of CD98 has not been linked to improved functionality. Nonetheless, it may represent an advantage in an amino acid-depleted environment such as the tumor microenvironment (TME), where there is a competition for nutrients between tumor cells, myeloid-derived suppressor cells and lymphocytes ^36,47^. Likewise, experiments in which glucose has been substituted with galactose revealed that glycolytic rates and effector functions of IL-2/12-or IL-12/15-stimulated NK cells decreased ^30,33^, thus highlighting the relevance of glucose availability. Indeed, in a mouse model, NK cells from lung cancer microenvironment showed lower levels of glycolysis and reduced effector functions ^35^. Moreover, the authors found that inhibition of FBP1 partially restored glycolytic activity and effector functions of lung NK cells ^35^. Although there are multiple factors that could modulate glycolysis and functionality of NK cells in the TME ^36^, an increased GLUT1 expression may mitigate the tumor-induced decrease of glycolytic activity due to a higher glucose uptake and thus be beneficial for NK cell functions. Therefore, a higher CD98 and GLUT1 expression could be helpful to support the functionality of CIML NK cells in nutrient-restricted environments.

Current knowledge indicates that there are several differences between non-preactivated and CIML NK cells at phenotypic and functional levels ^11,12,23,25–27,48,49,13–16,19–22^. Considering that metabolism supports cellular functions, we hypothesized that CIML NK cells may also differ in their metabolic profile. Others have previously reported that NK cell stimulation induced an increase in the glycolytic activity ^30,32,33,37,39,43,46,50^. In accordance, our results showed that NK cells have a higher glycolysis levels following IL-12/15/18-stimulation for 16-18 hours. Remarkably, we found that the increase in the glycolytic machinery is not a transient event, and that CIML NK cells retain elevated glycolytic activity for at least one week after IL-12/15/18-preactivation. It would be interesting to study if other stimuli could also induce a sustained metabolic reprogramming. For instance, it has been reported that adaptive NK cells from patients infected with cytomegalovirus (CMV) showed higher glycolytic metabolism than canonical NK cells ^51^, suggesting that viral infection could induce this glycolytic profile. Nonetheless, adaptive and CIML NK cells differ in other metabolic features. Adaptive NK cells from CMV+ individuals showed increased OXPHOS activity and higher SRC than canonical NK cells ^51^. IL-12/15/18-preactivated NK cells share the increment in OXPHOS and SRC rates following cytokine-stimulation, but these parameters return to basal levels after seven days. Nevertheless, in our experimental settings, CIML NK cells are not exposed to activating stimuli (e.g. viral infected cells) during the culture period of seven days, which may explain why OXPHOS activity returns to basal levels. Furthermore, mitochondrial mass of mouse NK cells did not show significant changes one week after CMV infection, and then decreased during the following days in the contraction-to-memory phase ^52^. Contrary to adaptive NK cells, CIML NK cells showed a progressive increase in the mitochondrial mass. Hence, adaptive and CIML NK cells exhibit different metabolic reprogramming. Nonetheless, considering the differences in the mitochondrial content of control and CIML NK cells, it would be interesting to study in future works if CIML NK cells are prone to increase more rapidly OXPHOS activity than canonical NK cells when restimulated after a resting period.

Our experiments were performed in a system with optimal conditions for NK cells. However, in the tumor context NK cells have to face a suppressive microenvironment that negatively affects to both metabolism and effector functions ^36^. In a recent report, control and CIML NK cells were cultured for seven days with ascites supernatant from patients with ovarian cancer, and then their effector functions were measured by co-culturing with ovarian cancer cell lines. Interestingly, the authors found that IL-12/15/18-preactivated NK cells showed a higher IFNγ production than control NK cells, suggesting that CIML NK cells may have the ability to overcome the suppressive soluble microenvironment ^13^. To further explore this idea, we decided to study effector functions of IL-12/15/18-preactivated NK cells in metabolically restricting conditions. Considering that NK cells showed reduced glycolytic activity in the TME ^35^ and to assess the relevance of this metabolic pathway for NK cell functions, we stimulated NK cells with either target cells or cytokines in the presence of the glycolysis inhibitor 2-DG. Previous works have shown that low doses (1 mM) of this metabolic inhibitor reduced IFNγ and/or granzyme B production of murine NK cells in vitro following stimulation with IL-15, IL-2/12 or anti-NK1.1 plus IL-2 ^32,33,43^. Remarkably, in vivo administration of 2-DG reduced IFNγ production but not TNF production of NK cells following poly(I:C)-mediated activation ^33^. Human NK cells are more resistant to 2-DG than murine NK cells ^32^, so direct comparisons should be carefully considered. Nonetheless, our data confirmed that distinct effector functions of human NK cells also have different sensitivity to 2-DG-induced glycolytic inhibition.

Interestingly, our results revealed that CIML NK cells restimulated with IL-12/15/18 at day 7 showed reduced TNF production and degranulation in the presence of 2-DG. Contrarily, those effector functions were not affected by 2-DG when NK cells were co-cultured with the K562 tumor cell line. It may be possible that K562 target cells hijacked 2-DG from media and thus NK cells were less exposed to this inhibitor. However, we have excluded this possibility because IFNγ and MIP-1β production were severely inhibited when 2-DG was added, so NK cells are also presumably exposed to this inhibitor. It has been previously described that IFNγ production is reduced by the combination of 2-DG and other metabolic inhibitors in murine NK cells stimulated through NK1.1, but no negative effects were found following IL-12/18 stimulation ^53^. Therefore, our results support the idea that metabolic requirements for a specific function could be different depending on the stimuli.

The effect of 2-DG on NK cell cytotoxic activity is still unclear. NK cells from CMV-infected mice receiving 2-DG treatment showed reduced target clearance ^32^. The same authors described that human NK cell cytotoxic activity against K562 cells was reduced when IL-15-stimulated NK cells were exposed to 2-DG for 24 hours prior to co-culturing NK and K562 cells ^32^. Contrarily, NK cells pretreated with 2-DG for 4 hours prior to the co-culture with K562 did not show defects in the killing of target cells, both in the presence and absence of 2-DG during the co-culture ^50^. Similar results were obtained by pretreating expanded human NK cells with 2-DG for 3 hours and co-culturing them with 721.221 target cells ^54^. However, the authors pointed that 2-DG-induced effect may be reversible, so its potential inhibition of NK cell-mediated cytotoxicity could be lost when the inhibitor is washed out ^54^. We assessed this issue by adding 2-DG during the co-culture period and we found that NK cell cytotoxicity against K562 cells is reduced by 2-DG. Discrepancies with previous studies could be explained because of the different experimental settings, such as co-culture duration, 2-DG concentration and different E:T ratios, among others. Importantly, it is known that NK cell target recognition involves an array of receptors that may be different depending on the nature of the target cell. This fact very possibly will have a relevant effect on the metabolic requirements of the effector functions of human NK cells, since, as we have shown, the requirements are stimuli-dependent. Moreover, it should be considered that in our experimental system K562 cells could also be affected by 2-DG, and that different target cells may be more or less dependent on glycolytic metabolism and thus have a higher or lower sensitivity to 2-DG. Therefore, the presence of 2-DG may differently affect to NK cell cytotoxicity against other cell lines, such as 721.221 cells (data not shown). Remarkably, our results showed that after IL-12/15/18-preactivation, CIML NK cells showed higher cytotoxic activity than control NK cells even in the presence of 2-DG. These results suggest that CIML NK cells could maintain higher antitumor activity in glycolysis-restricting conditions, such as the TME ^36^. It would be interesting to further explore whether CIML NK cells could overcome metabolic suppression, and understand the contribution of other metabolic pathways to their increased effector functions.

Immunometabolism could greatly influence the outcome of cancer immunotherapies. This idea has gained acceptance during the last years and, consequently, several strategies have been proposed to induce metabolic reprogramming of immune cells both in vivo and in vitro ^55^. Our work has demonstrated that IL-12/15/18 stimulation induces some metabolic changes that are maintained when cells return to a resting state, thus revealing that human CIML NK cells show durable metabolic reprogramming. Furthermore, we have explored the different glycolytic requirements for the effector functions of NK cells. These findings would help to design future therapies in which NK cells could be metabolically reprogrammed prior to cell infusion, and highlight the necessity to continue exploring the impact of TME over NK cell metabolism to find new means of improving NK cell-based cancer immunotherapy.

## ACKNOWLEDGMENTS

The authors thank the Cell Analytics Facility from the Achucarro Basque Center for Neuroscience (Leioa, Spain). The authors acknowledge A. Caro-Maldonado for technical assistance with extracellular flux assays. This work was supported by the following grants: AECC-Spanish Association Against Cancer (PROYE16074BORR) and Health Department, Basque Government (2019222027). I.T. is recipient of a fellowship from the Jesús de Gangoiti Barrera Foundation (FJGB18/002) and a predoctoral contract funded by the Department of Education, Basque Government (PRE_2019_2_0109). A.O. is recipient of a fellowship from the Jesús de Gangoiti Barrera Foundation (FJGB19/002). O.Z. is recipient of a postdoctoral contract funded by “Instituto de Salud Carlos III-Contratos Sara Borrell 2017 (CD17/00128)” and the European Social Fund (ESF)-The ESF invests in your future. F.B. is an Ikerbasque Research Professor, Ikerbasque, Basque Foundation for Science.

## AUTHOR CONTRIBUTIONS

F.B., O.Z. and I.T. designed the experiments; I.T. and A.M. performed the experiments; I.T., A.O. and A.M. analyzed the data; I.T., A.O., J.V., O.Z. and F.B. contributed to the interpretation of the data; I.T. drafted the manuscript and all authors edited the manuscript.

## REFERENCES

1 Zhang C, Hu Y, Shi C. Targeting Natural Killer Cells for Tumor Immunotherapy. Front Immunol 2020; 11. doi: 10.3389/fimmu.2020.00060.

2 Tarazona R, Lopez-Sejas N, Guerrero B, Hassouneh F, Valhondo I, Pera A et al. Current progress in NK cell biology and NK cell-based cancer immunotherapy. Cancer Immunol Immunother 2020. doi: 10.1007/s00262-020-02532-9.

3 Terrén I, Orrantia A, Mikelez-Alonso I, Vitallé J, Zenarruzabeitia O, Borrego F. NK Cell-Based Immunotherapy in Renal Cell Carcinoma. Cancers (Basel) 2020; 12: 316.

4 Mehta RS, Rezvani K. Chimeric Antigen Receptor Expressing Natural Killer Cells for the Immunotherapy of Cancer. Front Immunol 2018; 9: 1–12.

5 Conlon KC, Lugli E, Welles HC, Rosenberg SA, Fojo AT, Morris JC et al. Redistribution, hyperproliferation, activation of natural killer cells and CD8 T cells, and cytokine production during first-in-human clinical trial of recombinant human interleukin-15 in patients with cancer. J Clin Oncol 2015; 33: 74–82.

6 Cooley S, He F, Bachanova V, Vercellotti GM, DeFor TE, Curtsinger JM et al. First-in-human trial of rhIL-15 and haploidentical natural killer cell therapy for advanced acute myeloid leukemia. Blood Adv 2019; 3: 1970–1980.

7 Romee R, Cooley S, Berrien-Elliott MM, Westervelt P, Verneris MR, Wagner JE et al. First-in-human phase 1 clinical study of the IL-15 superagonist complex ALT-803 to treat relapse after transplantation. Blood 2018; 131: 2515–2527.

8 Margolin K, Morishima C, Velcheti V, Miller JS, Lee SM, Silk AW et al. Phase I trial of ALT-803, a novel recombinant IL15 complex, in patients with advanced solid tumors. Clin Cancer Res 2018; 24: 5552–5561.

9 Pérez-Martínez A, Fernández L, Valentín J, Martínez-Romera I, Corral MD, Ramírez M et al. A phase I/II trial of interleukin-15-stimulated natural killer cell infusion after haplo-identical stem cell transplantation for pediatric refractory solid tumors. Cytotherapy 2015; 17: 1594–1603.

10 Vela M, Corral D, Carrasco P, Fernández L, Valentín J, González B et al. Haploidentical IL-15/41BBL activated and expanded natural killer cell infusion therapy after salvage chemotherapy in children with relapsed and refractory leukemia. Cancer Lett 2018; 422: 107–117.

11 Romee R, Rosario M, Berrien-Elliott MM, Wagner JA, Jewell BA, Schappe T et al. Cytokine-induced memory-like natural killer cells exhibit enhanced responses against myeloid leukemia. Sci Transl Med 2016; 8: 357ra123.

12 Ni J, Miller M, Stojanovic A, Garbi N, Cerwenka A. Sustained effector function of IL-12/15/18–preactivated NK cells against established tumors. J Exp Med 2012; 209: 2351–2365.

13 Uppendahl LD, Felices M, Bendzick L, Ryan C, Kodal B, Hinderlie P et al. Cytokine-induced memory-like natural killer cells have enhanced function, proliferation, and in vivo expansion against ovarian cancer cells. Gynecol Oncol 2019; 153: 149–157.

14 Zhuang L, Fulton RJ, Rettman P, Sayan AE, Coad J, Al-Shamkhani A et al. Activity of IL-12/15/18 primed natural killer cells against hepatocellular carcinoma. Hepatol Int 2019; 13: 75–83.

15 Bonanni V, Antonangeli F, Santoni A, Bernardini G. Targeting of CXCR3 improves anti-myeloma efficacy of adoptively transferred activated natural killer cells. J Immunother Cancer 2019; 7: 1–16.

16 Boieri M, Ulvmoen A, Sudworth A, Lendrem C, Collin M, Dickinson AM et al. IL-12, IL-15, and IL-18 pre-activated NK cells target resistant T cell acute lymphoblastic leukemia and delay leukemia development in vivo. Oncoimmunology 2017; 6: 1–12.

17 Gang M, Marin ND, Wong P, Neal CC, Marsala L, Foster M et al. CAR-modified memory-like NK cells exhibit potent responses to NK-resistant lymphomas. Blood 2020. doi: 10.1182/blood.2020006619.

18 Stary V, Stary G. NK Cell-Mediated Recall Responses: Memory-Like, Adaptive, or Antigen-Specific? Front Cell Infect Microbiol 2020; 10: 1–7.

19 Cooper MA, Elliott JM, Keyel PA, Yang L, Carrero JA, Yokoyama WM. Cytokine-induced memory-like natural killer cells. Proc Natl Acad Sci U S A 2009; 106: 1915–9.

20 Terrén I, Mikelez I, Odriozola I, Gredilla A, González J, Orrantia A et al. Implication of Interleukin-12/15/18 and Ruxolitinib in the Phenotype, Proliferation, and Polyfunctionality of Human Cytokine-Preactivated Natural Killer Cells. Front Immunol 2018; 9: 737.

21 Wagner JA, Berrien-Elliott MM, Rosario M, Leong JW, Jewell BA, Schappe T et al. Cytokine-Induced Memory-Like Differentiation Enhances Unlicensed Natural Killer Cell Antileukemia and FcγRIIIa-Triggered Responses. Biol Blood Marrow Transplant 2017; 23: 398–404.

22 Leong JW, Chase JM, Romee R, Schneider SE, Sullivan RP, Cooper MA et al. Preactivation with IL-12, IL-15, and IL-18 Induces CD25 and a Functional High-Affinity IL-2 Receptor on Human Cytokine-Induced Memory-like Natural Killer Cells. Biol Blood Marrow Transplant 2014; 20: 463–473.

23 Ghofrani J, Lucar O, Dugan H, Reeves RK, Jost S. Semaphorin 7A modulates cytokine-induced memory-like responses by human natural killer cells. Eur J Immunol 2019; 49: 1153–1166.

24 Romee R, Schneider SE, Leong JW, Chase JM, Keppel CR, Sullivan RP et al. Cytokine activation induces human memory-like NK cells. Blood 2012; 120: 4751–4760.

25 Ewen E-M, Pahl JHW, Miller M, Watzl C, Cerwenka A. KIR downregulation by IL-12/15/18 unleashes human NK cells from KIR/HLA-I inhibition and enhances killing of tumor cells. Eur J Immunol 2018; 48: 355–365.

26 Lusty E, Poznanski SM, Kwofie K, Mandur TS, Lee DA, Ashkar AA. IL-18/IL-15/IL-12 synergy induces elevated and prolonged IFN-γ production by ex vivo expanded NK cells which is not due to enhanced STAT4 activation. Mol Immunol 2017; 88: 138–147.

27 Vendrame E, Fukuyama J, Strauss-Albee DM, Holmes S, Blish CA. Mass Cytometry Analytical Approaches Reveal Cytokine-Induced Changes in Natural Killer Cells. Cytom Part B-Clin Cytom 2017; 92: 57–67.

28 Kobayashi T, Mattarollo SR. Natural killer cell metabolism. Mol Immunol 2017;: 0–1.

29 Gardiner CM. NK cell metabolism. J Leukoc Biol 2019; 105: 1235–1242.

30 Keating SE, Zaiatz-Bittencourt V, Loftus RM, Keane C, Brennan K, Finlay DK et al. Metabolic Reprogramming Supports IFN-γ Production by CD56 bright NK Cells. J Immunol 2016; 196: 2552–2560.

31 O’Brien KL, Finlay DK. Immunometabolism and natural killer cell responses. Nat Rev Immunol 2019; 19: 282–290.

32 Mah AY, Rashidi A, Keppel MP, Saucier N, Moore EK, Alinger JB et al. Glycolytic requirement for NK cell cytotoxicity and cytomegalovirus control. JCI Insight 2017; 2: 1–17.

33 Donnelly RP, Loftus RM, Keating SE, Liou KT, Biron CA, Gardiner CM et al. mTORC1-Dependent Metabolic Reprogramming Is a Prerequisite for NK Cell Effector Function. J Immunol 2014; 193: 4477–4484.

34 Michelet X, Dyck L, Hogan A, Loftus RM, Duquette D, Wei K et al. Metabolic reprogramming of natural killer cells in obesity limits antitumor responses. Nat Immunol 2018; 19: 1330–1340.

35 Cong J, Wang X, Zheng X, Wang D, Fu B, Sun R et al. Dysfunction of Natural Killer Cells by FBP1-Induced Inhibition of Glycolysis during Lung Cancer Progression. Cell Metab 2018; 28: 243–255.e5.

36 Terrén I, Orrantia A, Vitallé J, Zenarruzabeitia O, Borrego F. NK Cell Metabolism and Tumor Microenvironment. Front Immunol 2019; 10: 1–9.

37 Velásquez SY, Himmelhan BS, Kassner N, Coulibaly A, Schulte J, Brohm K et al. Innate Cytokine Induced Early Release of IFNγ and CC Chemokines from Hypoxic Human NK Cells Is Independent of Glucose. Cells 2020; 9: 734.

38 Loftus RM, Assmann N, Kedia-Mehta N, O’Brien KL, Garcia A, Gillespie C et al. Amino acid-dependent cMyc expression is essential for NK cell metabolic and functional responses in mice. Nat Commun 2018; 9: 2341.

39 Almutairi SM, Ali AK, He W, Yang DS, Ghorbani P, Wang L et al. Interleukin-18 up-regulates amino acid transporters and facilitates amino acid–induced mTORC1 activation in natural killer cells. J Biol Chem 2019; 294: 4644–4655.

40 Salzberger W, Martrus G, Bachmann K, Goebels H, Heß L, Koch M et al. Tissue-resident NK cells differ in their expression profile of the nutrient transporters Glut1, CD98 and CD71. PLoS One 2018; 13: e0201170.

41 Zaiatz-Bittencourt V, Finlay DK, Gardiner CM. Canonical TGF-β Signaling Pathway Represses Human NK Cell Metabolism. J Immunol 2018; 200: 3934–3941.

42 Jensen H, Potempa M, Gotthardt D, Lanier LL. Cutting Edge: IL-2–Induced Expression of the Amino Acid Transporters SLC1A5 and CD98 Is a Prerequisite for NKG2D-Mediated Activation of Human NK Cells. J Immunol 2017; 199: 1967–1972.

43 Kedia-Mehta N, Choi C, McCrudden A, Littwitz-Salomon E, Fox PG, Gardiner CM et al. Natural Killer Cells Integrate Signals Received from Tumour Interactions and IL2 to Induce Robust and Prolonged Anti-Tumour and Metabolic Responses. Immunometabolism 2019; 1: e190014.

44 Nicholas D, Proctor EA, Raval FM, Ip BC, Habib C, Ritou E et al. Advances in the quantification of mitochondrial function in primary human immune cells through extracellular flux analysis. PLoS One 2017; 12: 1–19.

45 Berthe A, Zaffino M, Muller C, Foulquier F, Houdou M, Schulz C et al. Protein N-glycosylation alteration and glycolysis inhibition both contribute to the antiproliferative action of 2-deoxyglucose in breast cancer cells. Breast Cancer Res Treat 2018; 171: 581–591.

46 Marçais A, Cherfils-Vicini J, Viant C, Degouve S, Viel S, Fenis A et al. The metabolic checkpoint kinase mTOR is essential for IL-15 signaling during the development and activation of NK cells. Nat Immunol 2014; 15: 749–757.

47 Chang C, Qiu J, O’Sullivan D, Buck MD, Noguchi T, Curtis JD et al. Metabolic Competition in the Tumor Microenvironment Is a Driver of Cancer Progression. Cell 2015; 162: 1229–1241.

48 Song Y, Hu B, Liu Y, Jin Z, Zhang Y, Lin D et al. IL-12/IL-18-preactivated donor NK cells enhance GVL effects and mitigate GvHD after allogeneic hematopoietic stem cell transplantation. Eur J Immunol 2018; 48: 670–682.

49 Huber CM, Doisne JM, Colucci F. IL-12/15/18-preactivated NK cells suppress GvHD in a mouse model of mismatched hematopoietic cell transplantation. Eur J Immunol 2015;: 1727–1735.

50 Wang Z, Guan D, Wang S, Chai LYA, Xu S, Lam K-P. Glycolysis and Oxidative Phosphorylation Play Critical Roles in Natural Killer Cell Receptor-Mediated Natural Killer Cell Functions. Front Immunol 2020; 11: 1–17.

51 Cichocki F, Wu C-Y, Zhang B, Felices M, Tesi B, Tuininga K et al. ARID5B regulates metabolic programming in human adaptive NK cells. J Exp Med 2018; 215: 2379–2395.

52 O’Sullivan TE, Johnson LR, Kang HH, Sun JC. BNIP3-and BNIP3L-Mediated Mitophagy Promotes the Generation of Natural Killer Cell Memory. Immunity 2015; 43: 331–342.

53 Keppel MP, Saucier N, Mah AY, Vogel TP, Cooper MA. Activation-Specific Metabolic Requirements for NK Cell IFN-γ Production. J Immunol 2015; 194: 1954–1962.

54 Schafer JR, Salzillo TC, Chakravarti N, Kararoudi MN, Trikha P, Foltz JA et al. Education-dependent activation of glycolysis promotes the cytolytic potency of licensed human natural killer cells. J Allergy Clin Immunol 2019; 143: 346–358.e6.

55 Guerra L, Bonetti L, Brenner D. Metabolic Modulation of Immunity: A New Concept in Cancer Immunotherapy. Cell Rep 2020; 32: 107848.

