## Supplemental Figures for "Human cytokine-induced memory-like NK cells preserve increased glycolysis but the glycolytic-dependence of their effector functions differ between stimuli"

### Figure S1

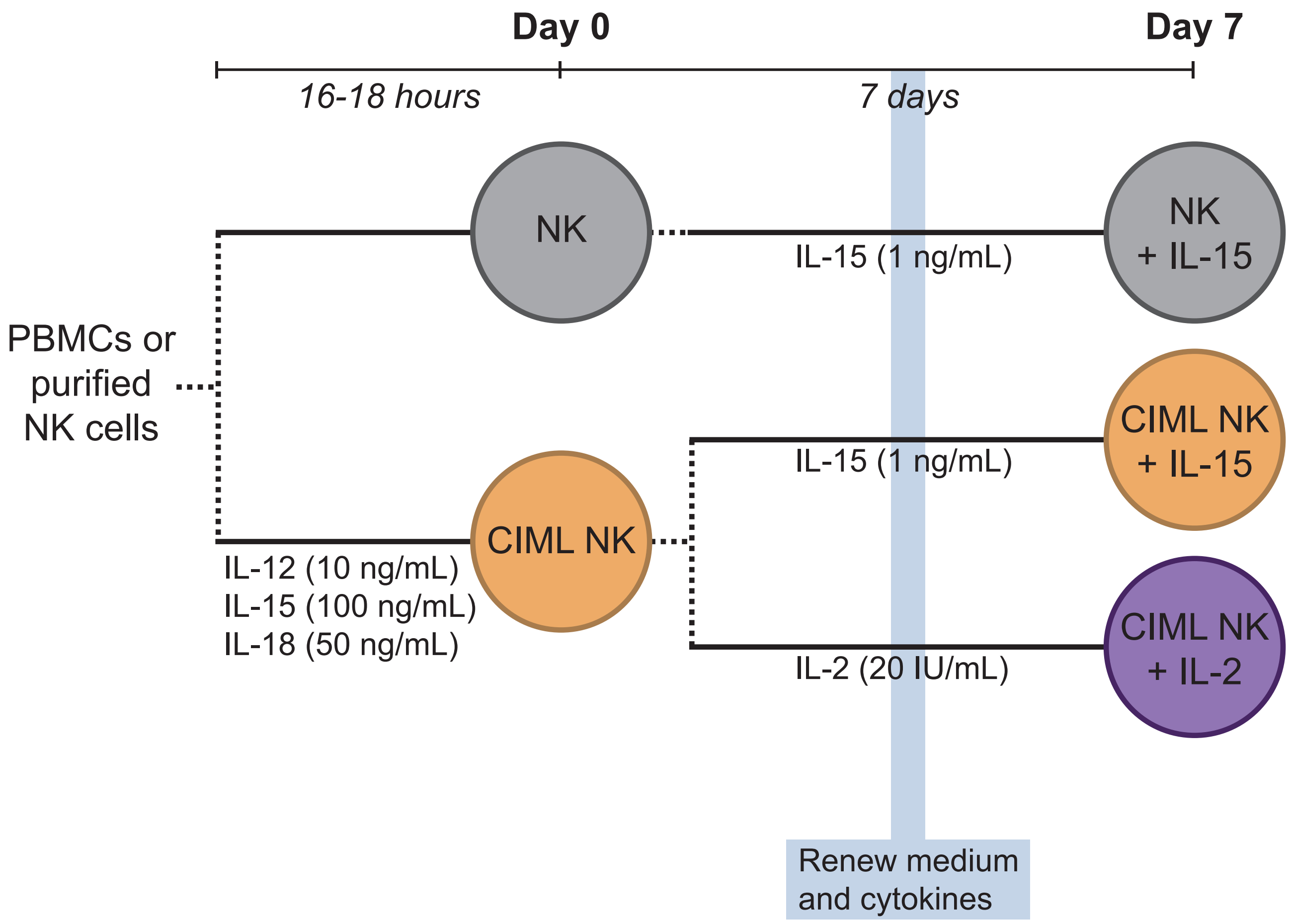


**Figure S1. Schematic representation of experimental design.** PBMCs or purified NK cells (depending on the assay, as indicated in the methodology section) were cultured for 16-18 hours with media alone (NK) or with a mixture of IL-12, IL-15 and IL-18 (10, 100 and 50 ng/mL, respectively) (CIML NK). At this time point (Day 0), cells were collected, washed and analyzed. Additionally, cells were further cultured for seven days in media containing 1 ng/mL IL-15 or 20 IU/mL IL-2. Media and cytokines were renewed four days after the Day 0. After seven days (Day 7), cells were collected, washed and analyzed.

### Figure S2

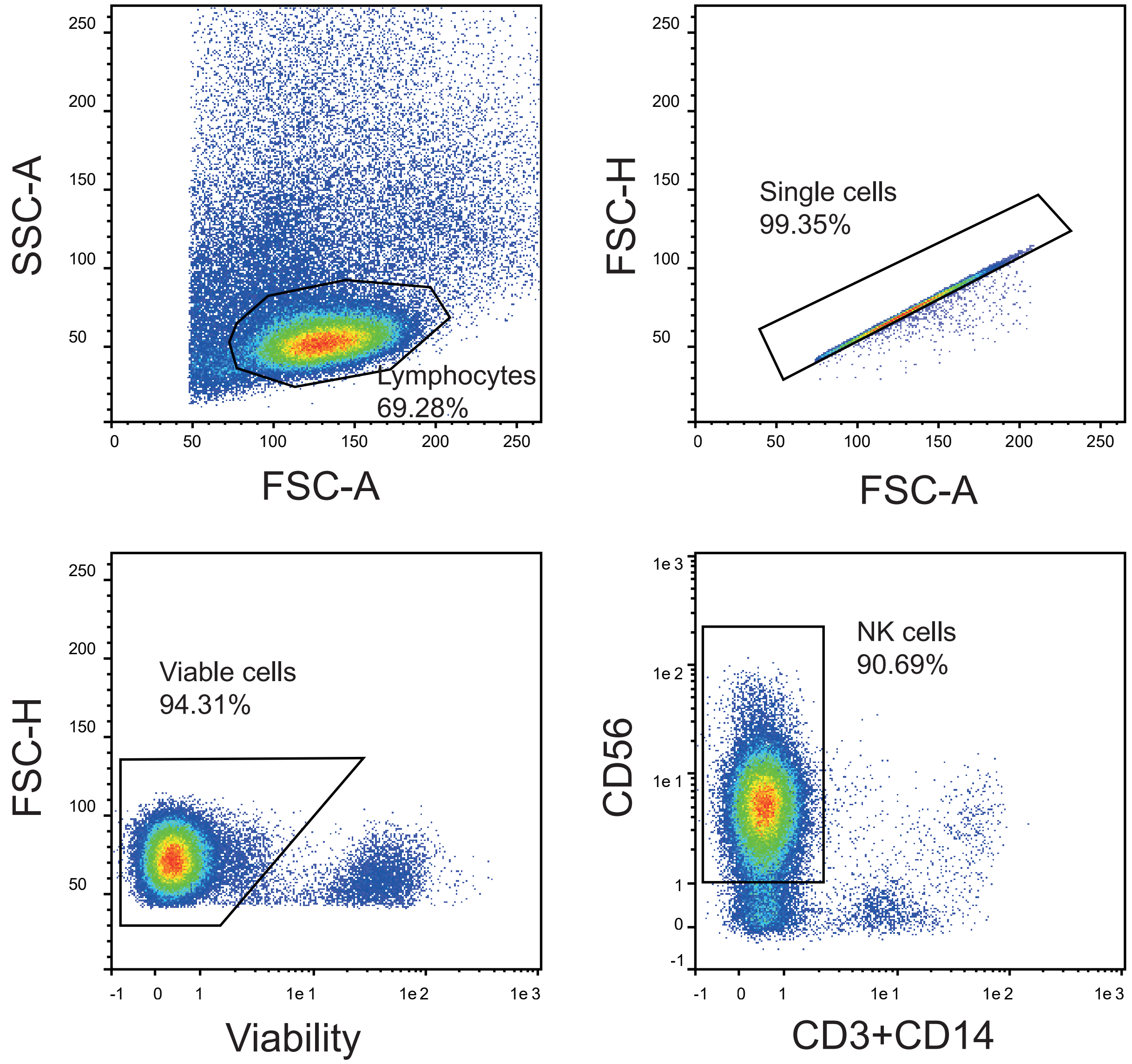


**Figure S2. Gating strategy of purified NK cells.** Lymphocytes were gated attending to forward and side scatter (FSC-A vs. SSC-A) parameters. Then, single cells were identified based on forward scatter (FSC-A vs. FSC-H), and dead cells were excluded with LIVE/DEAD Fixable Near-IR Dead Cell Stain Kit. Finally, NK cells were gated as CD3-CD14- and CD56+.

### Figure S3

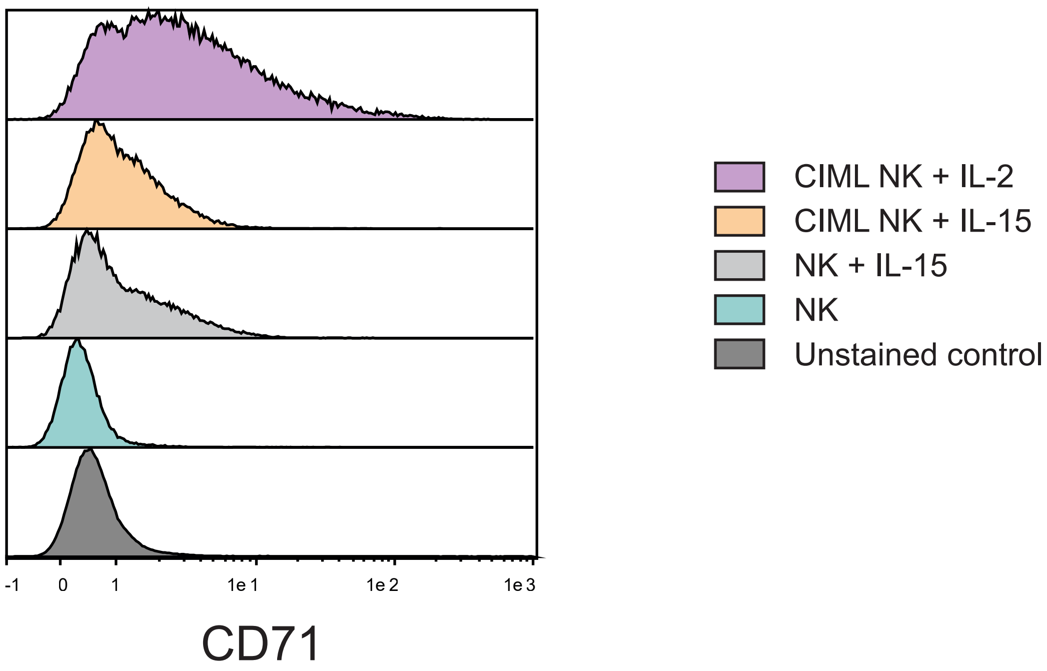


**Figure S3. Effect of low doses of IL-15 on CD71 expression.** Control (NK) and CIML NK cells were cultured in media without cytokines, or with IL-15 (1 ng/mL) or IL-2 (20 IU/mL) for seven days. Media was replaced during this culture period, as explained in the methodology section. Histograms show the expression of transferrin receptor CD71, measured within viable NK cells.

Figure S4

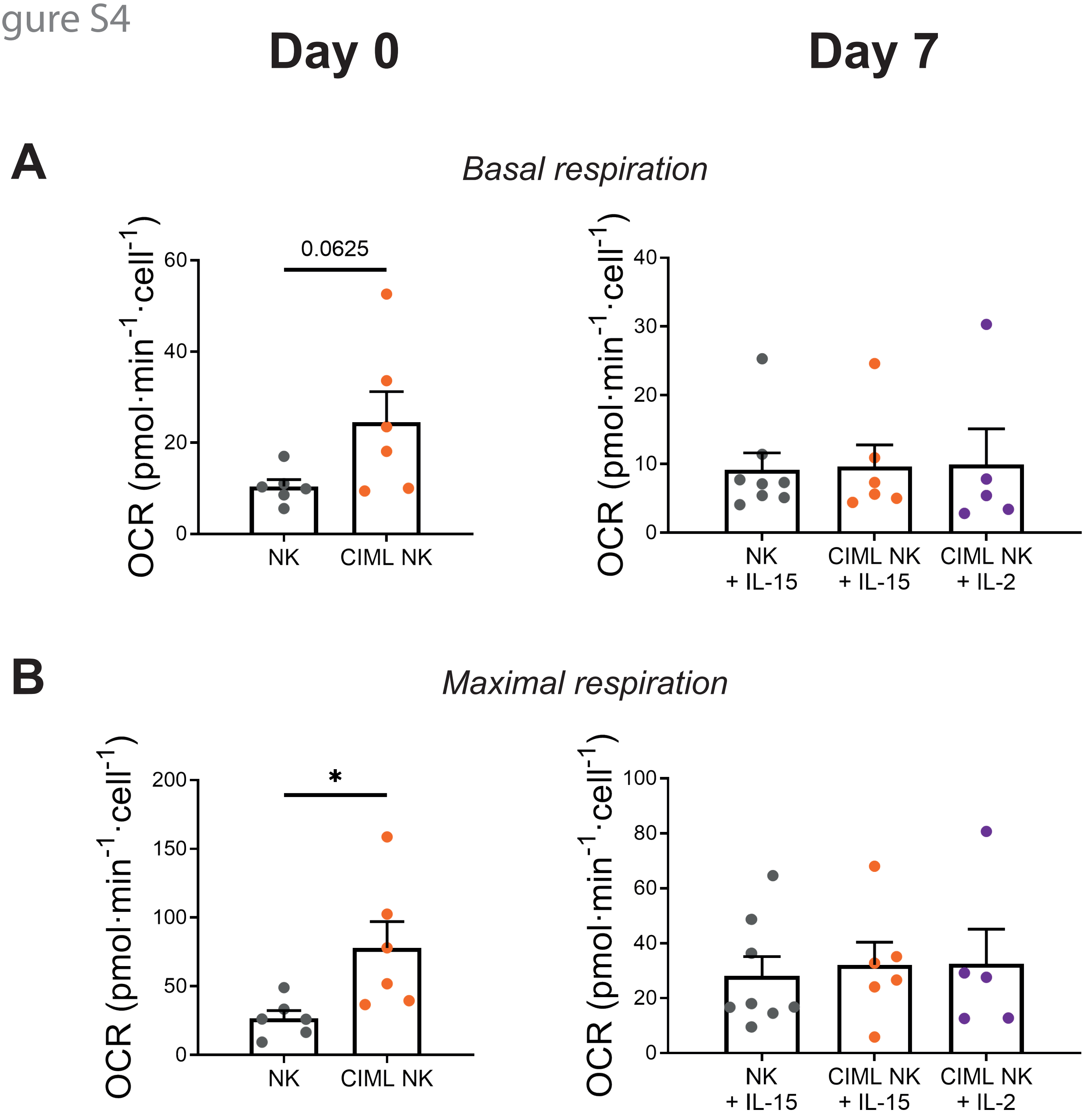


**Figure S4. Mitochondrial respiration of human cytokine-induced memory-like (CIML) NK cells.** Bar charts representing (A) basal respiration and (B) maximal respiration (i.e. respiration levels following addition of FCCP) rates, measured as oxygen consumption rate (OCR). Graphs show data of control NK cells (NK) and IL-12/15/18-stimulated NK cells (CIML NK) after 16-18 hours (Day 0, left column), and after 7 days of culture with IL-15 (NK+IL-15 or CIML NK+IL-15) or IL-2 (CIML NK+IL-2) (Day 7, right column). Means ± SEM are depicted. Statistical analyses were performed using Wilcoxon matched-pairs signed rank test. Each dot represents an independent experiment from a different donor (n = 5-8). $^*p<0.05$.

Figure S5

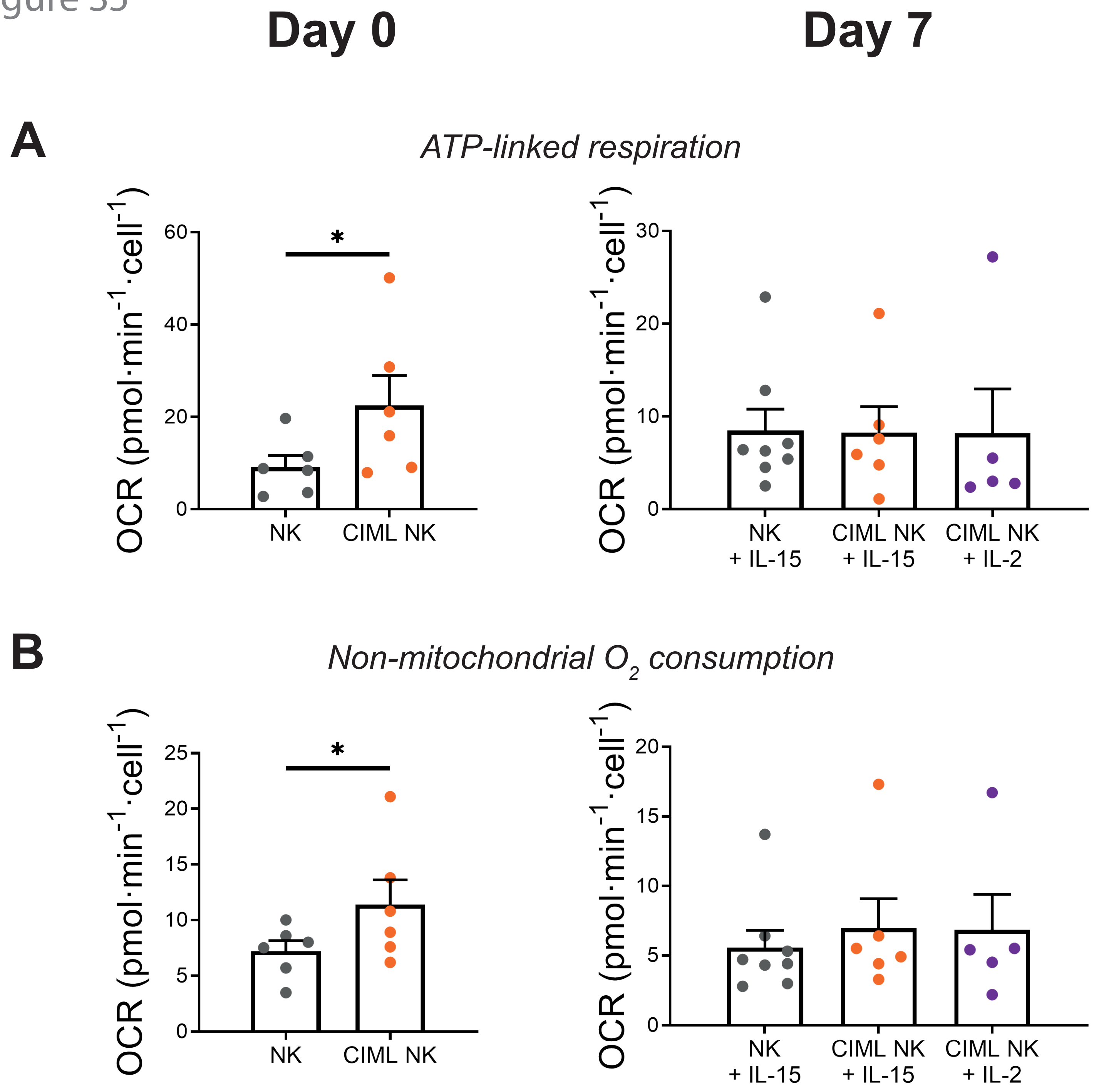


**Figure S5. Mitochondrial respiration-linked parameters of human cytokine-induced memory-like (CIML) NK cells.** Bar charts representing (A) ATP-linked respiration, measured by the decrease in oxygen consumption rate (OCR) when cells are exposed to ATP synthase inhibitor oligomycin, and (B) non-mitochondrial oxygen consumption, measured as the OCR of cells exposed to oligomycin, FCCP, rotenone and antimycin A. Graphs show data of control NK cells (NK) and IL-12/15/18-stimulated NK cells (CIML NK) after 16-18 hours (Day 0, left column), and after 7 days of culture with IL-15 (NK+IL-15 or CIML NK+IL-15) or IL-2 (CIML NK+IL-2) (Day 7, right column). Means ± SEM are depicted. Statistical analyses were performed using Wilcoxon matched-pairs signed rank test. Each dot represents an independent experiment from a different donor (n = 5-8). *$p<0.05$.

Figure S6

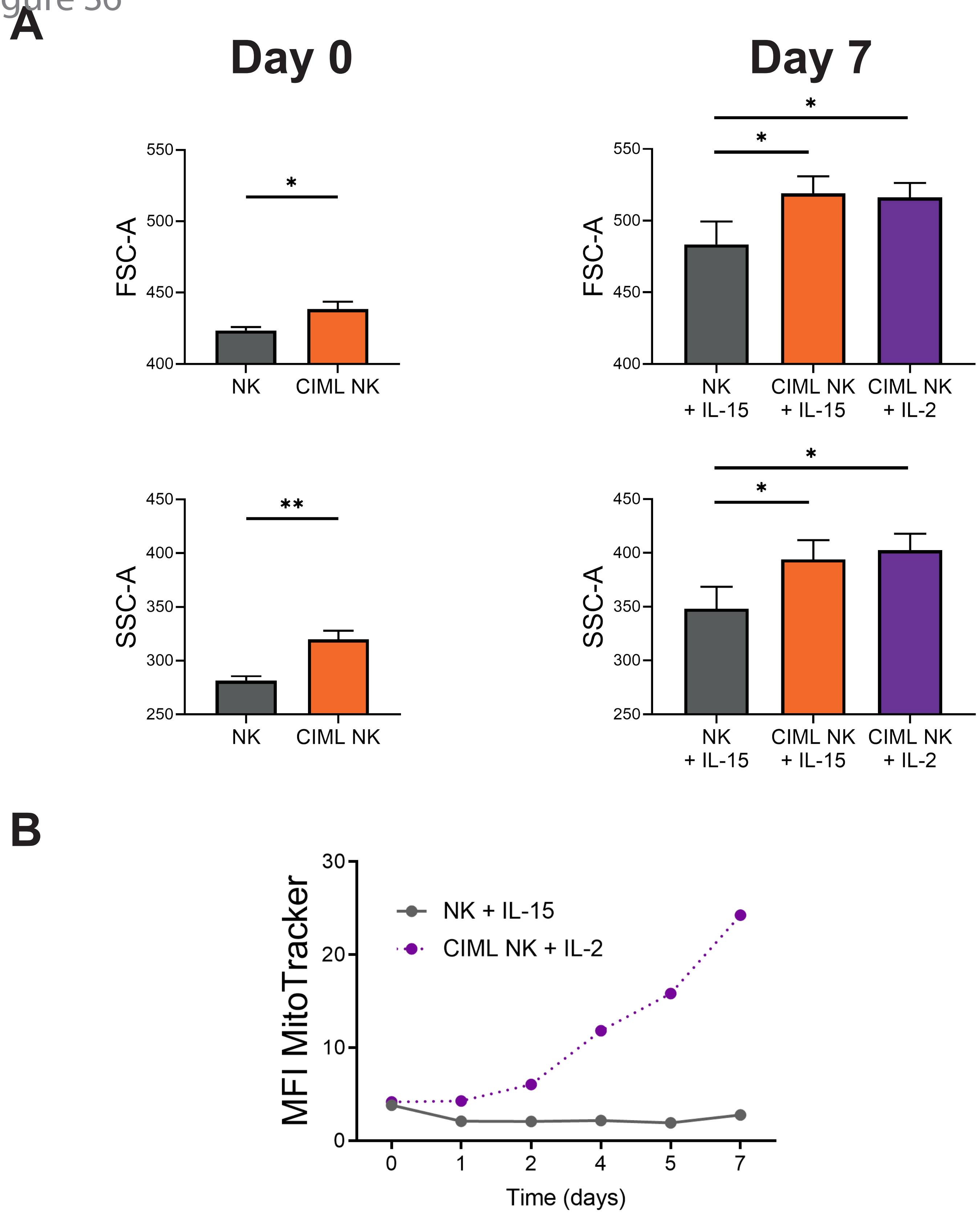


**Figure S6. Evolution of mitochondrial mass and cellular size of human cytokine-induced memory-like (CIML) NK cells.** (A) Bar graphs representing cellular size and granularity, measured by forward (FSC) and side scatter (SSC) values, of control NK cells (NK) and IL-12/15/18-stimulated NK cells (CIML NK) after 16-18 hours (Day 0, left column), and after 7 days of culture with IL-15 (NK+IL-15 or CIML NK+IL-15) or IL-2 (CIML NK+IL-2) (Day 7, right column). Means ± SEM are depicted. Statistical analyses were performed using Wilcoxon matched-pairs signed rank test (n = 8-9). *p<0.05, **p<0.01. (B) Repeated measures of mitochondrial mass during the culture period of seven days, measured as the median fluorescence intensity (MFI) of MitoTracker Green (n = 1).

### Figure S7

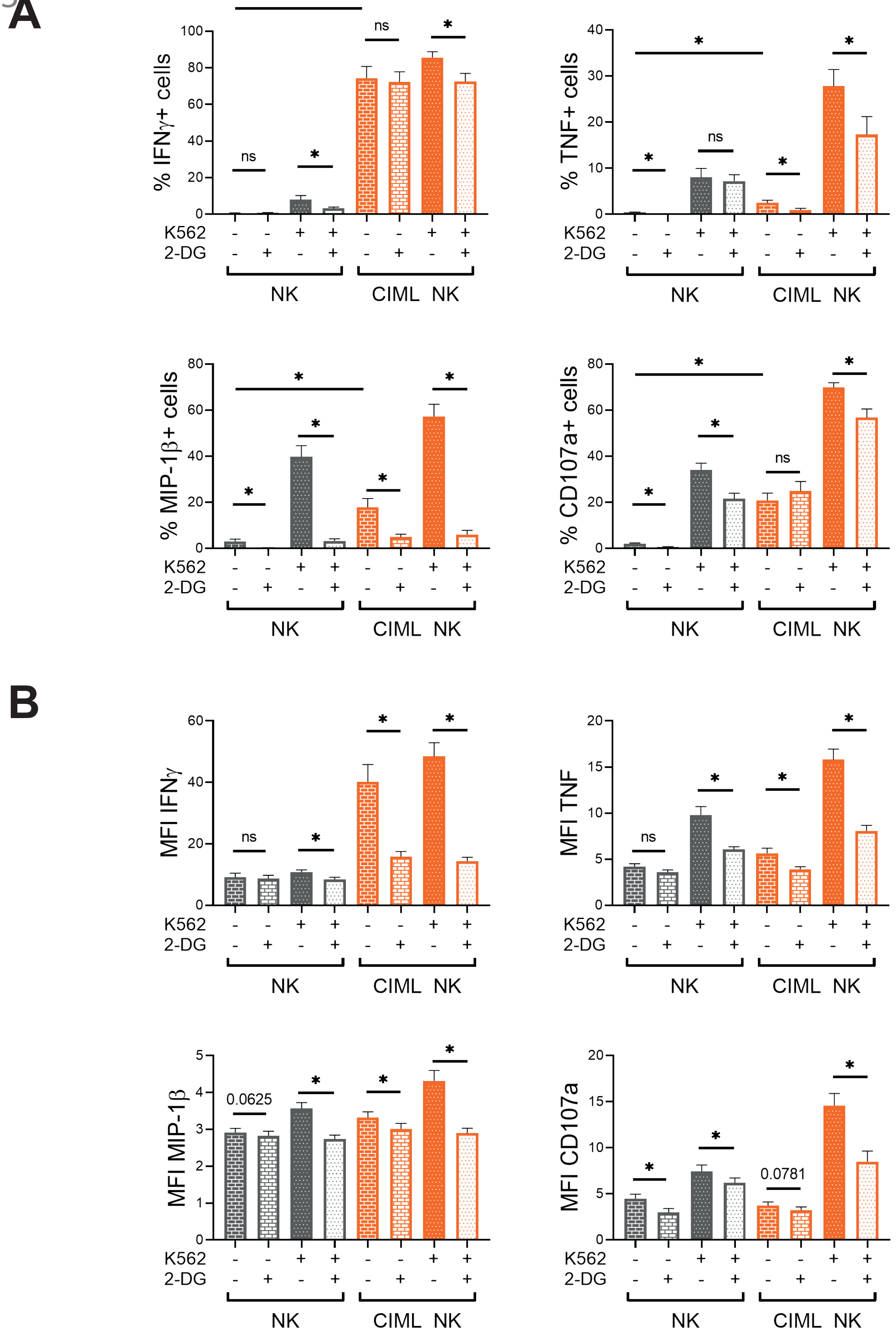

A
% IFNγ+ cells
% TNF+ cells
% MIP-1β+ cells
% CD107a+ cells
B
MFI IFNγ
MFI TNF
MFI MIP-1β
MFI CD107a
0.0625
0.0781
ns
*
K562
2-DG
NK
CIML NK

### Figure S7

**Figure S7. 2-DG-induced inhibition of cytokine/chemokine production and degranulation of human cytokine-induced memory-like (CIML) NK cells at day 0.** Control (NK) and CIML NK cells were co-cultured with K562 target cells (E:T ratio = 1:1) for 7 hours in the presence and absence of 50 mM 2-DG. Bar graphs showing (A) percentage of positive cells and (B) median fluorescence intensity (MFI) of the cells that are positive for IFNγ, TNF and MIP-1β, or degranulate (CD107a). Means ± SEM are depicted. Statistical analyses were performed using Wilcoxon matched-pairs signed rank test (n = 7). ns = non-significant, *$p<0.05$.

Figure S8

A

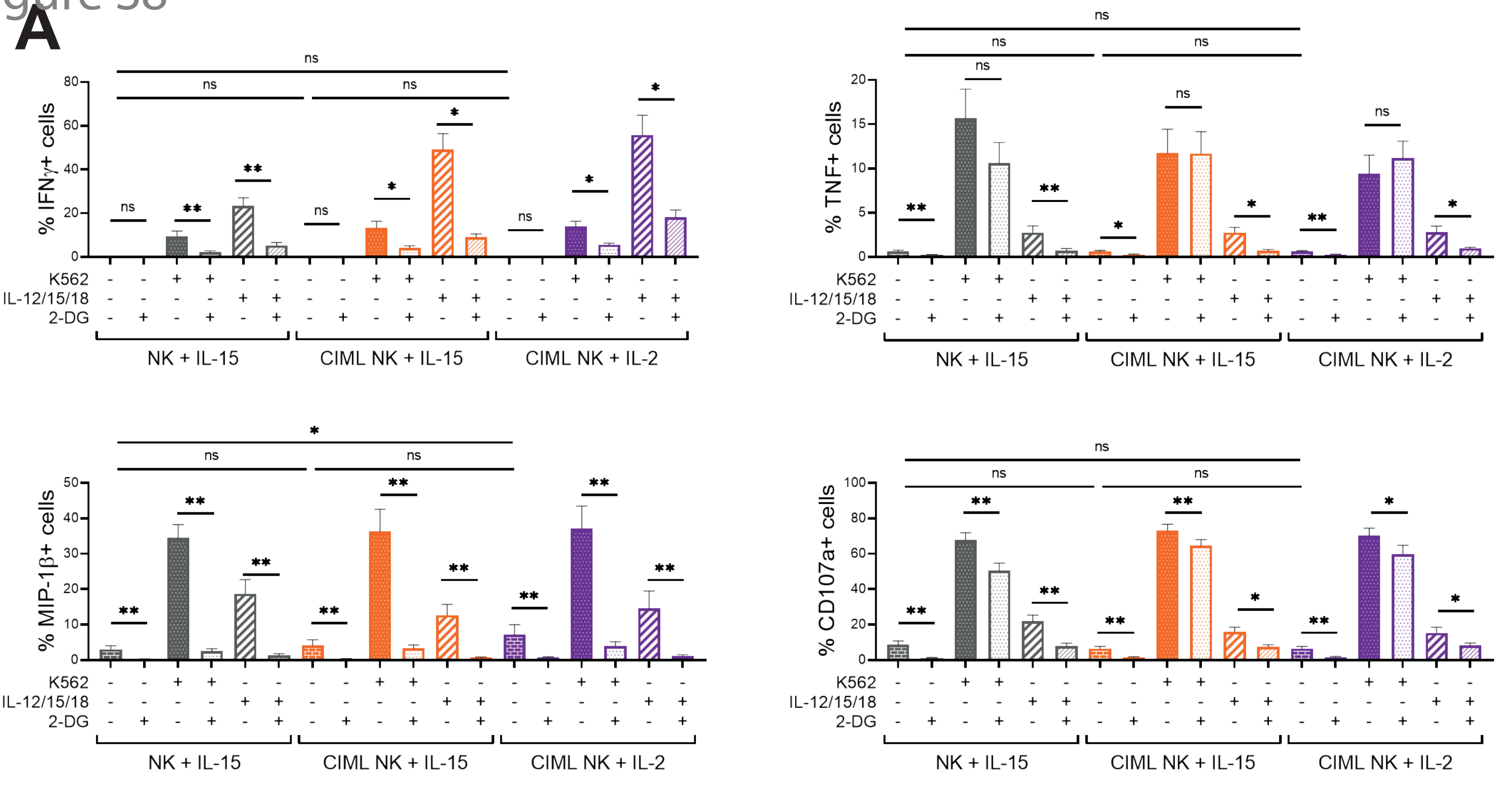
% IFNγ+ cells
% TNF+ cells
% MIP-1β+ cells
% CD107a+ cells
K562
IL-12/15/18
2-DG
NK + IL-15
CIML NK + IL-15
CIML NK + IL-2
ns

B

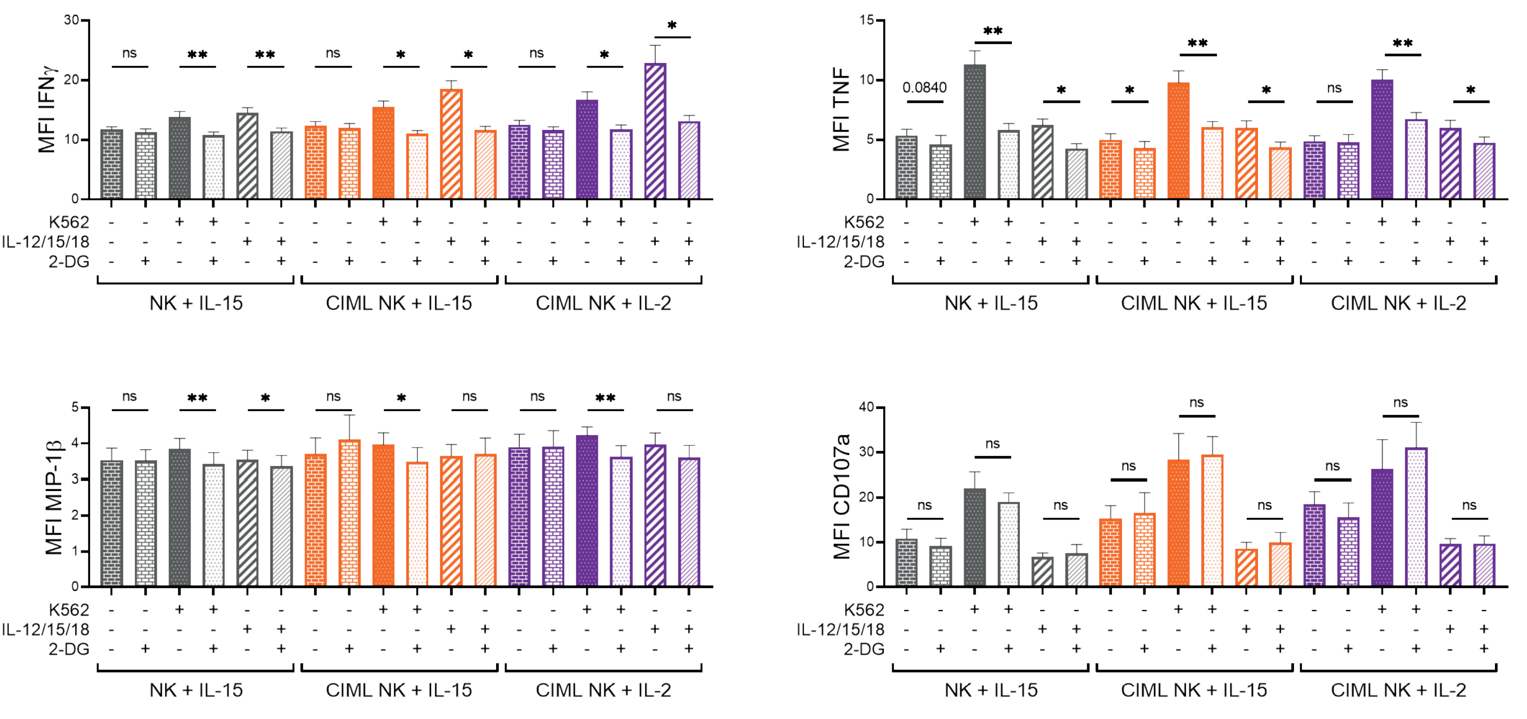
MFI IFNγ
MFI TNF
MFI MIP-1β
MFI CD107a
0.0840
K562
IL-12/15/18
2-DG
NK + IL-15
CIML NK + IL-15
CIML NK + IL-2
ns

### Figure S8

**Figure S8. 2-DG-induced inhibition of cytokine/chemokine production and degranulation of human cytokine-induced memory-like (CIML) NK cells at day 7.** Control (NK+IL-15) and CIML NK cells cultured for seven days with IL-15 (CIML NK+IL-15) or IL-2 (CIML NK+IL-2), were co-cultured with K562 target cells (E:T ratio = 1:1) or stimulated with IL-12, IL-15 and IL-18 (10, 100 and 50 ng/mL, respectively) for 7 hours in the presence and absence of 50 mM 2-DG. Bar graphs showing (A) percentage of positive cells and (B) median fluorescence intensity (MFI) of the cells that are positive for IFNγ, TNF and MIP-1β, or degranulate (CD107a). Means ± SEM are depicted. Statistical analyses were performed using Wilcoxon matched-pairs signed rank test (n = 7-10). ns = non-significant, *$p<0.05$, **$p<0.01$.
